# ERK signaling promotes resistance to TRK kinase inhibition in NTRK fusion-driven glioma mouse models

**DOI:** 10.1101/2024.03.13.584849

**Authors:** Sebastian Schmid, Zachary R Russell, Alex Shimura Yamashita, Madeline E West, Abigail G Parrish, Julia Walker, Dmytro Rudoy, James Z Yan, David C Quist, Betemariyam N Gessesse, Neriah Alvinez, Patrick J Cimino, Debra K Kumasaka, Ralph E Parchment, Eric C Holland, Frank Szulzewsky

**Affiliations:** Human Biology Division, Fred Hutchinson Cancer Center, Seattle, WA 98109, USA; Clinical Pharmacodynamic Biomarkers Program, Applied/Developmental Research Directorate, Frederick National Laboratory for Cancer Research, Leidos Biomedical Research, Inc., Frederick, MD 21701, USA; Surgical Neurology Branch, National Institute of Neurological Disorders and Stroke, National Institutes of Health, Bethesda, MD 20892, USA; Seattle Translational Tumor Research Center, Fred Hutchinson Cancer Center, Seattle, WA 98109, USA

**Author notes:** Contributed equally (Sebastian Schmid, Zachary R Russell). Correspondence: Frank Szulzewsky.

## Abstract

Pediatric-type high-grade gliomas frequently harbor gene fusions involving receptor tyrosine kinase genes, including neurotrophic tyrosine kinase receptor (NTRK) fusions. Clinically, these tumors show high initial response rates to tyrosine kinase inhibition but ultimately recur due to the accumulation of additional resistance-conferring mutations. Here, we developed a series of genetically engineered mouse models of treatment-naïve and –experienced NTRK1/2/3 fusion-driven gliomas. Both the TRK kinase domain and the N-terminal fusion partners influenced tumor histology and aggressiveness. Treatment with TRK kinase inhibitors significantly extended survival of NTRK fusion-driven glioma mice in a fusion– and inhibitor-dependent manner, but tumors ultimately recurred due to the presence of treatment-resistant persister cells. Finally, we show that ERK activation promotes resistance to TRK kinase inhibition and identify MEK inhibition as a potential combination therapy. These models will be invaluable tools for preclinical testing of novel inhibitors and to study the cellular responses of NTRK fusion-driven gliomas to therapy.

## Introduction

Tropomyosin receptor kinases (TRKA-C, encoded by the NTRK1-3 genes) belong to the family of receptor tyrosine kinases (RTKs), are activated by a series of ligands (NGF, BDNF and others), and are involved in neuronal development ^1^. The canonical TRK receptor variants consist of an extracellular ligand binding domain, a transmembrane domain, and an intracellular cytoplasmic kinase domain. Under physiological conditions, the expression and activity of these receptors is tightly regulated; ligand binding induces receptor dimer formation and subsequent autophosphorylation of several tyrosine residues in the activation loop of the receptor kinase domain. By contrast, in NTRK fusion proteins, a 5’ upstream partner that frequently induces dimerization is fused to the TRK tyrosine kinase domain, leading to its cytoplasmic localization as well as constitutive and de-regulated activation independent of ligand binding ^2^.

Gene fusions involving the three NTRK genes are estimated to be present in up to 1% of all solid tumors and occur with a multitude of different upstream fusion partners ^2, 3^. Large-scale next-generation sequencing efforts have identified recurrent NTRK gene fusions in subsets of pediatric-type high-grade gliomas ^4, 5, 6^ with a prevalence ranging from 0.4 to 40%, depending on the exact tumor type ^7^. In these pediatric-type tumors, the NTRK fusions are frequently the only known oncogenic drivers and potentially the tumor-initiating events. In addition to being found in pediatric-type gliomas, NTRK fusions are also found in adult-type gliomas, albeit at a lower frequency and frequently along with other common aberrations (such as gain of chromosome 7 and loss of chromosome 10), suggesting that they may not be the initiators or main oncogenic driver mutations in these tumors. Recurrent NTRK fusions have also been identified in a variety of peripheral cancers (including salivary-gland tumors, soft-tissue sarcomas, infantile fibrosarcoma, as well as thyroid, colon, skin, and lung cancers), both in pediatric and adult patients ^8^.

Clinical trials with the first-generation TRK-specific tyrosine kinase inhibitors (TKIs) Larotrectinib (LOXO-101) and Entrectinib (RXDX-101) have reported high overall response rates in NTRK fusion-positive solid tumors, including both peripheral tumors and brain tumors (both primary and metastatic) ^3, 8, 9, 10, 11^. However, these responses are transient, and tumors ultimately progress due to the presence of residual persister cells that ultimately acquire additional primary mutations in the kinase domain of the NTRK fusion or secondary mutations in downstream pathway effectors such as Ras or Raf and eventually lead to tumor recurrence. Although novel second generation TKIs, such as Seletrectinib (LOXO-195) and Repotrectinib (TPX-0005) can overcome some of these resistances, tumors frequently also become resistant to these inhibitors ^12, 13, 14^. The median duration of treatment with Larotrectinib for patients with glioma was 11.1 months, with around half of the patients discontinuing treatment due to progression of disease ^3^. The inability of current clinical inhibitors to elicit a long-term tumor regression highlights the necessity to better understand the biology of NTRK fusions and of fusion-positive tumors, as well as the cellular responses of these tumors to tyrosine kinase inhibition *in vivo*.

Genetically engineered mouse models (GEMMs) of human cancers are valuable tools for preclinical drug testing and for studying the underlying oncogenic drivers and activated molecular pathways in these tumors ^15, 16^. GEMMs have several advantages over traditional patient-derived xenograft models, including the use of immune-competent mice as well as the ability to introduce specific driver mutations in a defined genetic background. We have previously used the RCAS/tv-a system for somatic cell gene transfer system to study the functions of common genetic drivers in glioma (such as PDGF or loss of NF1) as well as gene fusions found in meningioma or ependymoma ^17, 18, 19, 20^. This system has specific advantages over germline GEMM models, such as the ability to rapidly modify or mutate the oncogenic driver or introduce additional drivers or tumor suppressor losses.

Although the genomic landscapes differ between adult-type and pediatric-type tumors, understanding what pathways are dysregulated by strong oncogenic drivers (such as gene fusions) in pediatric-type gliomas can be used to understand the overall biology of cancer and adult glioma. Several unanswered questions remain about the biology and oncogenic capacities of these fusions. Are these gene fusions sufficient to induce tumors experimentally in vivo and do the different fusions vary in their oncogenic potential? Do the different NTRK fusions respond to each TKI equally and does the therapeutic response seen in culture mimic that seen in vivo?

In this study, we addressed these questions by modeling eight different NTRK fusion-driven mouse gliomas using the RCAS/tv-a system. We found that all eight tested NTRK fusions were able to induce tumors when expressed in mice but observed significant differences in the oncogenic potentials of different fusions. The oncogenesis was significantly enhanced when introducing additional tumor suppressor losses, most importantly *Cdkn2a* loss. We then determined the therapeutic response of NTRK fusion-driven cell line models to TKIs in vitro and subsequently performed pre-clinical in vivo trials, treating mice harboring NTRK fusion-driven gliomas with the two first-generation TKIs Larotrectinib and Entrectinib. In vivo, we observed significantly increased survival rates for all fusion tumor types, however, all tumors ultimately recurred after treatment discontinuation due to the presence of treatment-resistant persister cells. In summary, our results suggest that NTRK gene fusions are strong oncogenic drivers and the likely tumor-initiating events in pediatric-type NTRK fusion-positive gliomas. Tyrosine kinase inhibition is able to significantly extend survival, however, ultimately failed to completely eliminate tumors, eventually resulting in tumor recurrence.

## Results

### Expression of NTRK1/2/3 fusions alone is sufficient for the formation of gliomas in mice

To determine if the expression of different *NTRK1/2/3* gene fusions is sufficient to induce the formation of glioma-like tumors in mice, we cloned the human coding sequences of nine *NTRK* fusions frequently found in human pediatric-type high-grade gliomas into the RCAS retroviral vector, including two *NTRK1* fusions (*TPM3-NTRK1* (*TN1*), *CHTOP-NTRK1* (*CN1*)), three *NTRK2* fusions (*GKAP1-NTRK2* (*GN2*), *NACC2-NTRK2* (*NN2*), and *QKI-NTRK2* (*QN2*)), and three *NTRK3* fusions (*ETV6-NTRK3* (*ETV6-N3*), *EML4-NTRK3* (*EML4-N3*), and *BTBD1-NTRK3* (*BN3*)) (Suppl. Figure S1A-B).

We used the RCAS/tv-a system for somatic cell gene transfer in combination with *Nestin*/tv-a (N/tv-a) mice ^16^, to intracranially express the different *NTRK* fusion constructs in Nestin-positive stem– and progenitor cells. While *NTRK* gene fusions are enriched in infantile/pediatric patients^4, 5, 6^, they are also found in adult patients ^21, 22^; we therefore intracranially expressed the different *NTRK* fusion constructs in mice of different ages (neonatal mice postnatal day 0-2 (p0-p2) or one-week-old mice (p7), as well as 5-7-week-old mice).

Each of the analyzed *NTRK* gene fusions was able to induce the formation of tumors from Nestin-expressing cells in the brain on their own without the loss of additional tumor suppressors (Suppl. Figure S1C). We observed prominent differences in tumor penetrance and aggressiveness depending on the age of transgene expression (Figure 1A-D, Suppl. Figure S1C-D). Tumors induced in neonatal mice were frequently of high penetrance and generally more aggressive, whereas tumors induced in 1-week-old mice displayed a high penetrance but were generally slow-growing and remained non-symptomatic even 120 days post-injection (study endpoint). In rare instances when tumors grew to a larger size and became symptomatic, these were usually centered around the third ventricle (Suppl. Figure S1E), suggesting that Nestin-positive cells in that area are more susceptible to oncogenic transformation by *NTRK* fusions compared to cells near the lateral ventricles at that point in development. Of note, *NTRK* gene fusions have been observed in human pediatric diffuse midline gliomas ^22^. Upon expression in adult *Cdkn2a* wild type mice, only *CHTOP-NTRK1* was able to induce the formation of small non-symptomatic tumors (2 out of 3 mice), whereas the other tested *NTRK* fusions failed to do so. In these adult *Cdkn2a* wild type tumors, only a minority of cells in the tumor stained positive for the TRK fusion kinase domain.

**Fig 1:**
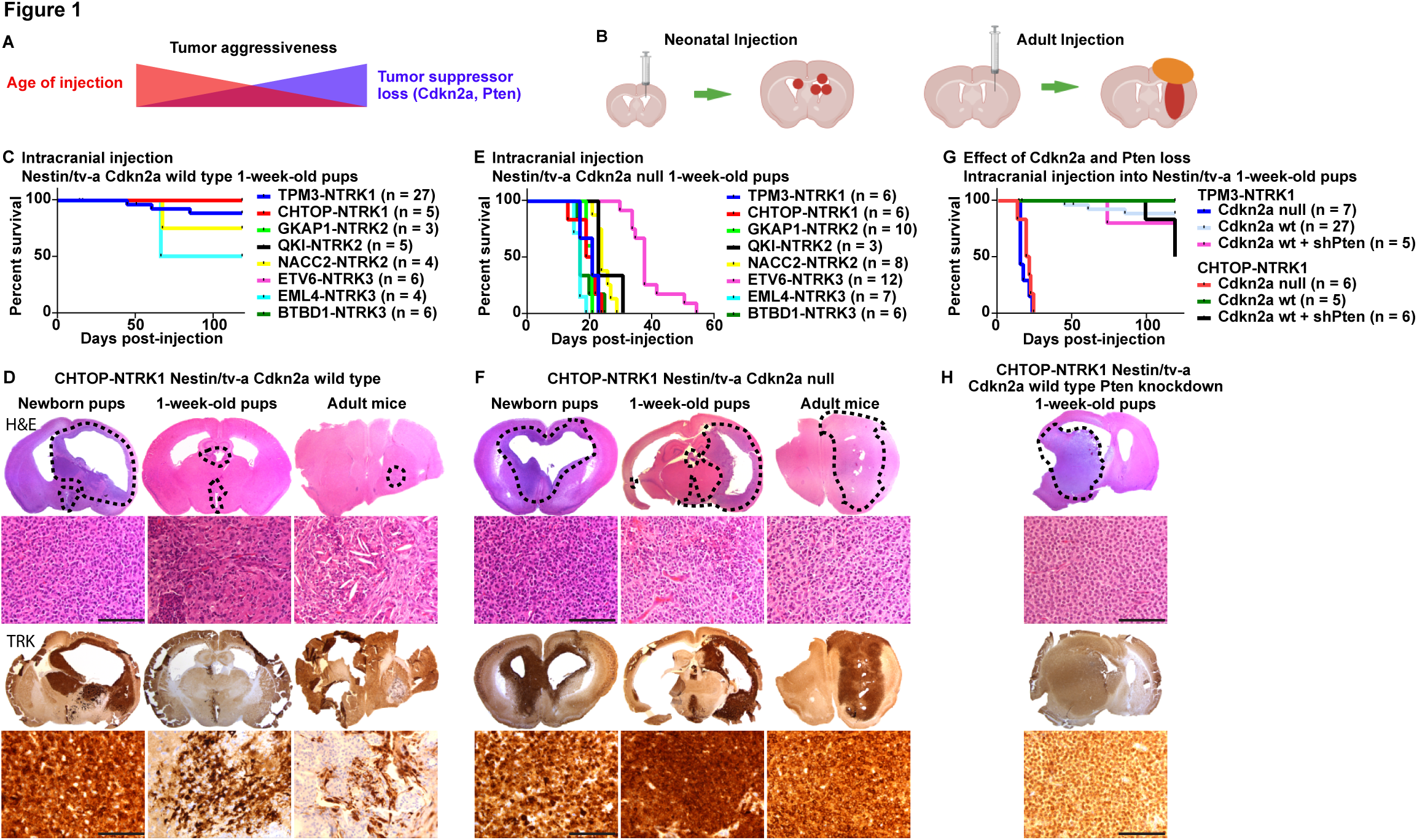
Expression of *NTRK* gene fusions in Nestin-positive cells induces the formation of experimental mouse gliomas. A) Aggressiveness of *NTRK* fusion-driven mouse gliomas is influenced by both mouse age at time of tumor induction and loss of additional tumor suppressors. B) Schematic depicting RCAS injection and tumor formation for neonatal and adult mice. C) Kaplan-Meier curve showing symptom-free survival of Nestin/tv-a *Cdkn2a* wild type mice upon intracranial expression of *NTRK* gene fusions. D) Representative stainings (H&E and IHC for the TRK kinase domain) of mouse gliomas induced by *CHTOP-NTRK1* in N/tv-a *Cdkn2a* wild type mice of different ages (Newborn pups, one-week-old pups, adult mice). Tumor aggressiveness decreases with increased age at tumor induction. E) Kaplan-Meier curve showing symptom-free survival of Nestin/tv-a *Cdkn2a* null mice upon intracranial expression of different *NTRK* gene fusion. F) Representative stainings (H&E and IHC for the TRK kinase domain) of mouse gliomas induced by expression of *CHTOP-NTRK1* in N/tv-a *Cdkn2a* null mice of different ages. G) Kaplan-Meier curve showing symptom-free survival of *Nestin*/tv-a *Cdkn2a* wild type mice upon intracranial expression of *TPM3-NTRK1* and *CHTOP-NTRK1* gene fusions with and without additional *Pten* knockdown. H) Representative stainings (H&E and IHC for the TRK kinase domain) of a mouse glioma induced by expression of *CHTOP-NTRK1* in N/tv-a *Cdkn2a* wild type mice with additional knockdown of *Pten*. Scale bar indicates 100 µm. Black dotted line indicates tumor boundaries (B,F,H).

Taken together, our data suggests that expression of *NTRK* fusions alone is sufficient to induce tumor formation without the loss of additional tumor suppressor and that these fusions are the likely tumor-initiating events in *NTRK* fusion-positive high-grade gliomas. The age of expression has significant influence on the aggressiveness of the resulting tumors.

### Additional loss of tumor suppressors (*Cdkn2a*, *Pten*) increases aggressiveness of *NTRK* fusion-driven gliomas

*CDKN2A/B* and *PTEN* loss are frequent events in adult-type IDH-wild type glioblastoma. Albeit at a lower frequency, deletions in *CDKN2A/B* and inactivating mutations in *PTEN* have also been reported in *NTRK1/2/3* fusion-positive pediatric-type high-grade gliomas ^4, 5, 6^. To assess if the loss of *Cdkn2a* affects the growth behavior of *NTRK* fusion-driven gliomas in our system, we intracranially expressed the different *NTRK* fusions in Nestin-positive cells of N/tv-a *Cdkn2a* null mice. In addition, we intracranially co-expressed *TN1* and *CN1* in combination with a short hairpin against *Pten* ^18^ in N/tv-a *Cdkn2a* wild type mice to model *Pten* loss.

Loss of *Cdkn2a* significantly enhanced tumor growth and increased tumor histologic grade (Figure 1E-F, Suppl. Figure S1F-H). The age at initial transgene expression significantly impacted tumor latency. Mice with tumors induced at p0 frequently became symptomatic before 21 days of age, however also caused hydrocephalus formation in a large percentage of mice, whereas tumors induced in adult mice showed a significantly longer latency compared to either p0 or p7 tumors. Adult-initiated tumors – regardless of the gene fusion – frequently developed an extra-axial and extra-cranial component, in addition to the intraparenchymal tumor component.

We observed significant differences in the latency of *Cdkn2a* null tumors derived from different *NTRK* fusions, in part even if they were derived from the same *NTRK1/2/3* variant (Figure 1E, Suppl. Figure S1F-H). Tumors induced by either *TN1*, *CN1*, *GN2*, *EML4-N3*, or *BN3* generally showed the lowest latency (17-21 days and 26-28 days median survival for p7 and adult tumors, respectively), follow by either *QN2* or *NN2* (23-24 days and 41-42 days median survival for p7 and adult tumors, respectively), whereas tumors induced by *ETV6-N3* displayed the longest latency (38 days and 53 days median survival for p7 and adult tumors, respectively). The *ETV6-N3* fusion sequence reported from glioma samples contains a breakpoint that generates a shorter kinase domain sequence compared to the other two *NTRK3* fusions in our cohort (Suppl Figure S1A-B). A variant of *ETV6-N3* with a longer kinase domain sequence – equal to *EML4-N3* and *BN3* – has been reported in other cancers (Suppl. Figure S1A-B) ^23^. Expression of this longer variant led to a non-significant trend (p = 0.09) towards longer latency (55 days versus 38 days median survival) (Suppl. Figure S1I). These results suggest that the longer latency of tumors induced by *ETV6-N3* compared to tumors induced by *EML4-N3* or *BN3* is not caused by the truncated *NTRK3* kinase domain present in *ETV6-N3*.

The majority of the *NTRK* gene fusion driven tumors (*TN1*, *CN1*, *GN2*, adult *NN2*, *ETV6-N3*, pediatric *QN2*) appeared histologically as diffuse astrocytic gliomas, demonstrating an infiltrating growth pattern and were of glial origin (Figure 1F, Suppl. Figure S1H, J). In general, these gliomas had variable mitotic activity and occasionally showed palisading necrosis. The astrocytic-appearing neoplastic cells in these tumors were characterized by enlarged, angulated, and hyperchromatic nuclei present on a fibrillary background. A couple of the *NTRK* fusion tumors (*EML4-N3*, pediatric *NN2*) appeared to have a more compact arrangement with the cells appearing more epithelioid, characterized by round cells with more defined cell borders, abundant eosinophilic cytoplasm, and variably enlarged and round nuclei with prominent nucleoli. One of the *NTRK* gene fusion tumors (adult *QN2*) had a predominantly spindled cell component, that was almost sarcomatous-like, and the cells had a compact fascicular arrangement.

Loss of *Pten* expression had a more subtle impact on tumor growth compared to Cdkn2a loss (Figure 1G-H, Suppl. Figure S1K). There was no significant difference in the survival of *TN1* or *CN1* tumors upon loss of *Pten* expression, however, a subset of tumors originating from the lateral ventricles were of larger size and became symptomatic. Histologically, these tumors often demonstrated oligodendroglial-like perinuclear clearing, which differed from that of *Pten* wild type or *Cdkn2a* null tumors ^24^.

As described above, adult-initiated tumors frequently consisted of an intra-parenchymal and a connected extra-parenchymal component. The extra-parenchymal component frequently grew as an extra-axial tumor and also grew out through the needle tract as extra-cranial tumor tissue. Similar to the intra-parenchymal tumor component, tumor cells in the extra-parenchymal tumors were also Olig2-positive, indicating a glial tumor-lineage (Suppl. Figure S1L). Magnetic resonance imaging (MRI) (T1-weighted and T2-weighted, pre– and post-contrast) on mice harboring intracranial *TPM3-NTRK1* tumors that exhibited extra-cranial tumor growth showed that the extra-cranial and to a lesser degree also the extra-axial tumor components were contrast enhancing, whereas the intraparenchymal tumor component was not. This suggests that the extra-cranial and extra-axial tumor components are more permeable to the contrast agent and therefore potentially also for small molecule inhibitors (Suppl. Figure S1M).

Since *NTRK* gene fusions are also found in non-CNS cancers (including lung, liver, and colon cancers), we also expressed these gene fusions in the flank and abdominal cavity of N/tv-a *Cdkn2a* null mice and observed the formation of highly aggressive sarcomatous-like spindle cell neoplasms that appeared to have arisen from the soft tissue and encased nearby organs such as colon and kidney (Suppl. Figure S1N-O).

Taken together, these results suggest that additional loss of tumor suppressors substantially increases the aggressiveness of *NTRK* fusion-driven gliomas. The N-terminal fusion partners can influence tumor aggressiveness and latency.

### Modeling of *NTRK* fusion-driven mouse gliomas harboring resistance-associated kinase domain point mutant variants

Targeted sequencing of recurrent patient tumors has recurrently identified point mutations in the kinase domain of *NTRK* gene fusion, suggesting that these mutations may constitute an early escape mechanism of *NTRK* fusion-positive tumor cells to bypass TKIs ^8^. The most common mutation hotspots include the gatekeeper residue (F589L in NTRK1), the solvent front residue (G595R in NTRK1), and xDFG mutations in the activation loop (G667C in NTRK1). The gatekeeper and the solvent front mutations are also frequently observed in combination ^2, 8^.

To assess if these kinase domain mutations influence tumor latency, we intracranially expressed the different *TPM3-NTRK1* kinase point mutant variants in one-week-old N/tv-a *Cdkn2a* null and wild type mice (Suppl. Figure S1Q-T). Similar to unmutated TPM3-NTRK1, all four mutant variants were able to rapidly induce tumor formation in a *Cdkn2a* null mice (Suppl. Figure S1R-S). The F589L, G595R, and F589L-G595R mutant variants displayed a latency similar to unmutated TPM3-NTRK1, whereas the G667C mutant version displayed a significantly longer latency (27 days versus 21 days, p = 0.0048).

In N/tv-a *Cdkn2a* wild type mice the F589L, G595R, and G667C mutant variants displayed a similar oncogenic potential compared to unmutated TPM3-NTRK1, with a high tumor penetrance, but tumors generally remained small and non-symptomatic even 120 days post-injection (Suppl. Figure S1P-Q), apart from a small number of symptomatic tumors originating from the third ventricle. By contrast, tumors induced by the expression of TPM3-NTRK1-F589L-G595R were significantly faster growing compared to tumors induced by unmutated TPM3-NTRK1 (p = 0.026), although the majority of tumors still remained non-symptomatic 120 days after induction. In addition, we observed the formation of large symptomatic tumors originating from the lateral ventricles in 2 out of 14 mice injected with TPM3-NTRK1-F589L-G595R (Suppl. Figure S1T), something not observed upon expression of any of the other constructs.

Taken together, our data suggests that by expressing *NTRK* fusion variants that harbor resistance-associated kinase point mutations, we can induce the formation of gliomas that mimic treatment-experienced tumors. Some of these kinase point mutations influence tumor aggressiveness.

### *NTRK* fusion-induced mouse gliomas show activation of the PI3K-AKT-S6 and the RAF-MEK-ERK pathways

To characterize these mouse gliomas in more detail, we performed immunohistochemistry (IHC) stainings for the TRK kinase domain, the glial markers Olig2 and GFAP, as well as the proliferative marker Ki67 (Figure 2, Suppl. Figure S2A-H). Tumors derived from *TN1*, *CN1*, *GN2*, *NN2*, or *ETV6-N3* were uniformly positive for the glial marker Olig2 regardless of the age of injection, whereas tumor cells frequently also stained positive for GFAP in pediatric, but not in adult tumors. Tumors derived from *QN2*, *EML4-N3*, or *BN3* fusions predominantly stained negative for GFAP (both pediatric and adult), and only a subset of tumor cells stained positive for Olig2. All tumors showed a high abundance of Ki67-positive cells.

**Fig 2:**
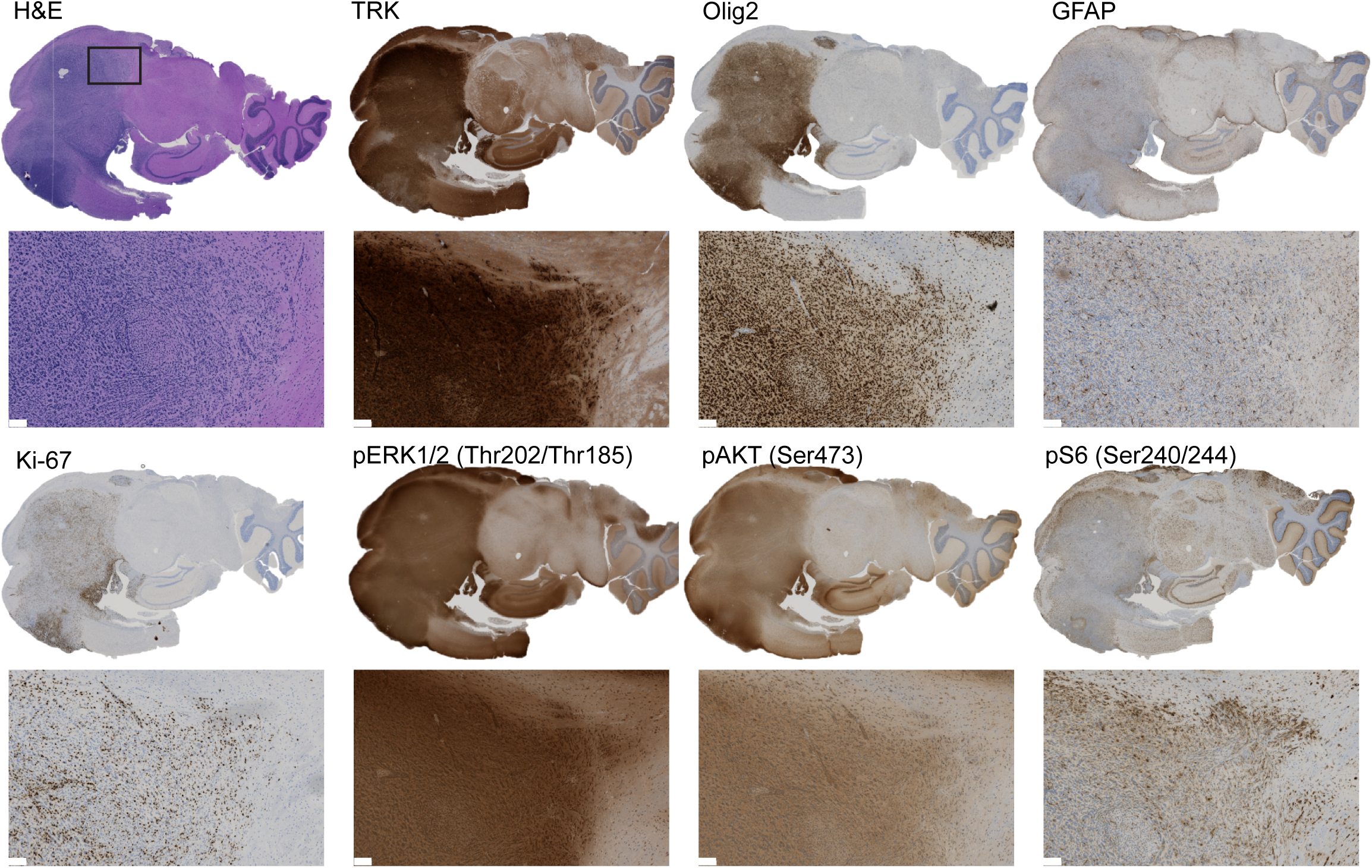
*NTRK* fusion-driven experimental mouse gliomas show activation of the PI3K-AKT-S6 and RAF-MEK-ERK pathways. Representative IHC stainings of a *TPM3-NTRK1*-driven mouse glioma for TRK (TRK kinase domain), Olig2, GFAP, Ki-67, phospho-ERK1/2 (Thr202/Thr185), phospho-AKT (Ser473), and phospho-S6 (Ser240/244). Scale bar indicates 100 µm.

Two of the main mitogenic downstream pathways activated by receptor tyrosine kinases, including NTRK receptors, are the PI3K-AKT-S6 and RAF-MEK-ERK pathways. To analyze if these pathways are also activated in the different NTRK fusion-driven mouse gliomas, we performed IHC stainings for phospho-ERK, phospho-AKT, and phospho-S6 and observed robust staining for all three epitopes in the different tumors, suggesting that both the PI3K-AKT– S6 and the RAF-MEK-ERK pathways are activated in these tumors (Figure 2, Suppl. Figure S2A-H).

Taken together, our results indicate that *NTRK* fusion-driven mouse tumors are of glial lineage and activate both the PI3K-AKT-S6 and the RAF-MEK-ERK pathway.

### Development of *NTRK* fusion cell line-based models to test the efficacy of various TKI against resistant-associated kinase point mutations in vitro

To assess the efficacy of TKI treatment against *NTRK* fusion-driven cells in vitro, we established several *NTRK* fusion-driven cell line models. We first transduced NIH-3T3-tv-a cells with RCAS viruses encoding for either GFP, TN1, GN2, or ETV6-N3 and subsequently performed 3D spheroid growth assays (Figure 3A, Suppl. Figure S3A-B). In the absence of a strong oncogenic driver NIH-3T3 cells are unable to grow in 3D spheroid conditions, whereas oncogenic signaling can induce the growth of these cell spheroids ^19, 20, 25^. While GFP-expressing cells did not grow as spheroids, spheroids derived from *NTRK* gene fusion-expressing cells continued to grow in size over several days (Figure 3B, Suppl Figure S3C). We tested the efficacy of four different TKIs – Larotrectinib, Entrectinib, as well as the two second-generation inhibitors Repotrectinib (TPX-0005) and Selitrectinib (LOXO-195) – to inhibit the growth of these spheroids (Figure 3C-E, Suppl. Figure S3C-E). Treatment with any of the four TKIs did not influence the growth of GFP-expressing spheroids, whereas the growth of *NTRK* fusion-driven spheroids was inhibited by all four TKIs in a dose-dependent manner. Entrectinib treatment was more effective in *TN1*– expressing cells, compared to cells expressing *GN2* or *ETV6-N3*, but there was otherwise no difference in the efficacy of the different TKIs between the different *NTRK* fusions (Suppl. Figure S3E).

**Fig 3:**
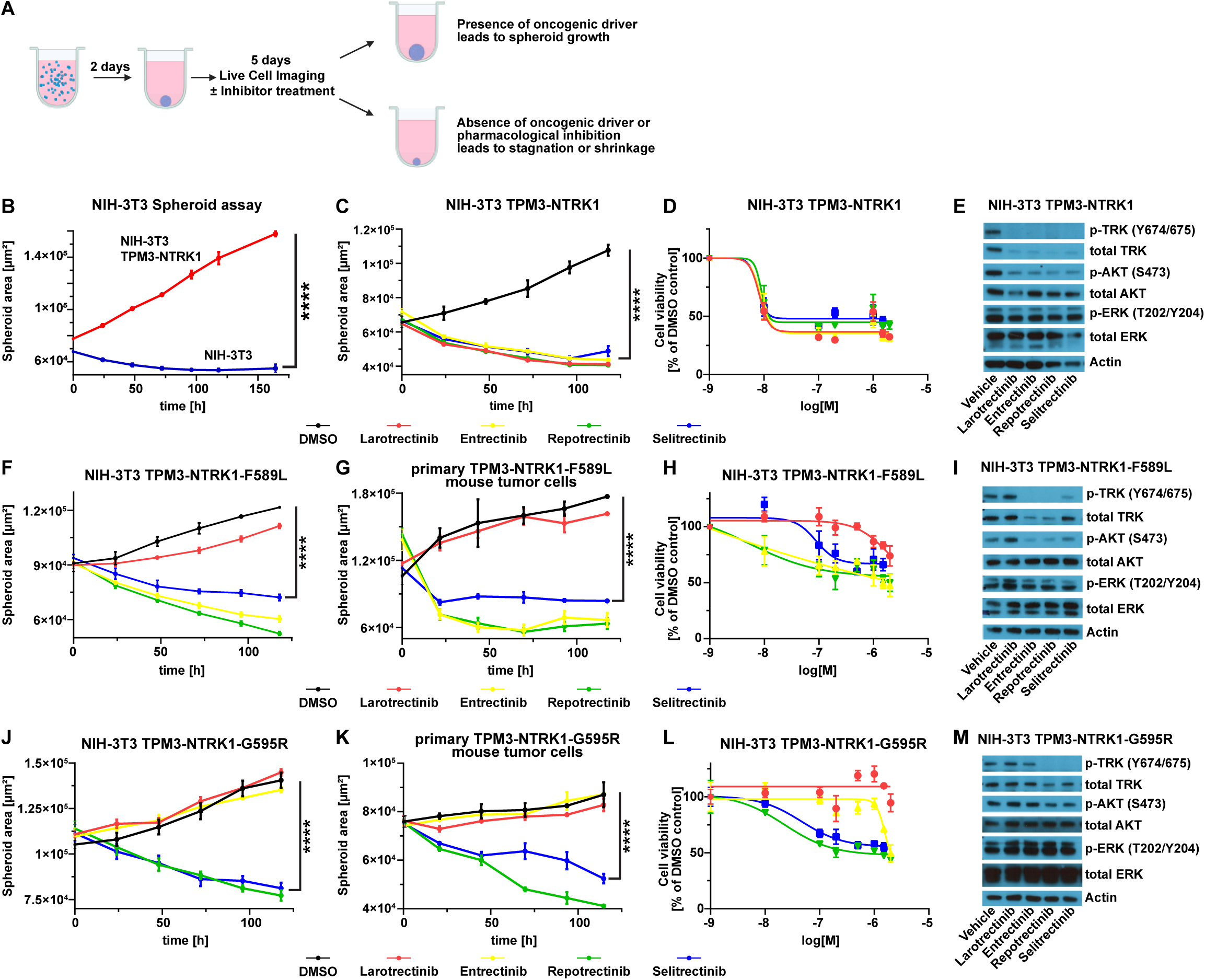
*NTRK* fusion-driven cells respond to TKI treatment in vitro. A) Schematic overview of the workflow of the 3D spheroid assay. B) Growth curve of a 3D spheroid assay with either untransduced NIH-3T3-tv-a cells (blue line) or cells expressing *TPM3-NTRK1* (red line). C) Growth curve of TPM3-NTRK1-expressing NIH-3T3-tv-a cell spheroids treated with either DMSO (0.1%) or indicated TKIs (100 nM each). D) Dose response curve of TPM3-NTRK1-expressing NIH-3T3-tv-a cell spheroids treated with different TKI concentrations (1, 10, 100, 200, 500 nM, 1 µM, 1.5 µM, or 2 µM) relative to DMSO-treated cells. E) Representative Western Blot of TPM3-NTRK1-expressing NIH-3T3-tv-a cells treated with 100 nM of indicated TKIs. F-G) Growth curve of TPM3-NTRK1-F589L-expressing NIH-3T3-tv-a (F) or primary mouse tumor (G) cell spheroids treated with either DMSO (0.1%) or indicated TKIs (100 nM each). H) Dose response curve of TPM3-NTRK1-F589L-expressing NIH-3T3-tv-a cell spheroids treated with different TKI concentrations (same as D) relative to DMSO-treated cells. I) Representative Western Blot of TPM3-NTRK1-F589L-expressing NIH-3T3-tv-a cells treated with 100 nM of indicated TKIs. J-K) Growth curve of TPM3-NTRK1-G595R-expressing NIH-3T3-tv-a (J) or primary mouse tumor (K) cell spheroids treated with either DMSO (0.1%) or indicated TKIs (100 nM each). L) Dose response curve of TPM3-NTRK1-G595R-expressing NIH-3T3-tv-a cell spheroids treated with different TKI concentrations (same as D) relative to DMSO-treated cells. M) Representative Western Blot of TPM3-NTRK1-G595R-expressing NIH-3T3-tv-a cells treated with 100 nM of indicated TKIs. Error bars show SEM. Analysis was done using ordinary Two-way ANOVA (B,C,F,G,J,K). (****) P < 0.0001.

Resistance to therapy against first-generation TKIs frequently occurs via the accumulation of *NTRK* kinase domain point mutations. To analyze how the different point mutations influence treatment response in vitro, we transduced NIH-3T3-tv-a cells with RCAS viruses encoding the different TPM3-NTRK1 mutants (F589L, G595R, G667C, F589L-G595R) and tested the efficacy of the different TKIs to inhibit the growth of these spheroids (Figure 3F-M. Suppl. Figure S3B,F-J). Cells expressing the F589L gatekeeper residue mutation were resistant to Larotrectinib, but still susceptible to Entrectinib, Repotrectinib, and Selitrectinib (Figure 3F, Suppl. Figure S3J), whereas the G595R solvent front mutation conferred resistance to both Larotrectinib and Entrectinib (Figure 3J, Suppl. Figure S3J). Cells expressing either the G667C xDFG mutation, or a combination of the F589L and G595R mutations were completely resistant to Selitrectinib and showed a decreased sensitivity to Repotrectinib (Suppl. Figure S3F-G, J). Western Blots of TKI-treated cells showed a decrease in p-TRK, p-ERK, and p-AKT levels that reflected the susceptibilities of the different resistance point mutations to the different TKIs (Figure 3E,I,M; Suppl Figure S3F-G).

We then established cell lines from primary mouse flank tumors induced by the injection of RCAS viruses encoding either TPM3-NTRK1-F589L, TPM3-NTRK1-G595R, or TPM3-NTRK1-F589L-G595R into N/tv-a *Cdkn2a* null mice and performed 3D spheroid assays with these cells. We again observed that TKI treatment was able to inhibit the growth of these spheroids in a resistance mutation-dependent manner that mirrored our findings from the NIH-3T3-tv-a cell line models (Figure 3G,K; Suppl. Figure S3H-I).

Taken together, *NTRK* fusion driven cell line models can predict the response of different kinase point mutants to first and second-generation TKIs.

### In vivo TKI treatment leads to the regression of *NTRK* gene fusion-driven mouse gliomas

To evaluate if TKIs can induce the regression of intracranial *NTRK* fusion-driven gliomas in vivo, we expressed an HA-tagged version of *EML4-N3* in N/tv-a *Cdkn2a* null mice and treated the resulting tumors with either Vehicle (two mice) or Entrectinib (60 mg/kg, two mice) twice-a-day for 24 hours (three injections in total). Mice were euthanized one hour after the third treatment, brains were formalin-fixed and subsequently analyzed by H&E and IHC.

Entrectinib treatment was able to induce the regression of the *EML4-N3*-driven mouse tumors after just 24 hours of treatment (Figure 4A-C). Entrectinib-treated tumors appeared smaller than Vehicle-treated tumors and displayed decreased levels of the proliferation marker phospho-Histone H3 and increased levels of apoptosis (Cleaved Caspase-3) (Figure A-B). Of note, Entrectinib treatment led to the decrease of total HA and Olig2 staining in a majority of tumor cells, suggesting that TKI treatment can induce the degradation of total NTRK fusion protein levels (Figure 4A). We observed a similar degradation of total TRK protein upon TKI treatment in vitro in both NIH-3T3 and primary mouse tumor cells (Figure 3E,I; Suppl. Figure S3D,I). In addition, we observed a prominent reduction in both p-ERK and p-S6 staining in Entrectinib-treated tumors. However, while a large percentage of tumor cells showed a decrease in these markers (HA, Olig2, p-ERK, p-S6), a subset of cells did not, suggesting that they may be less affected by TKI treatment.

**Fig 4:**
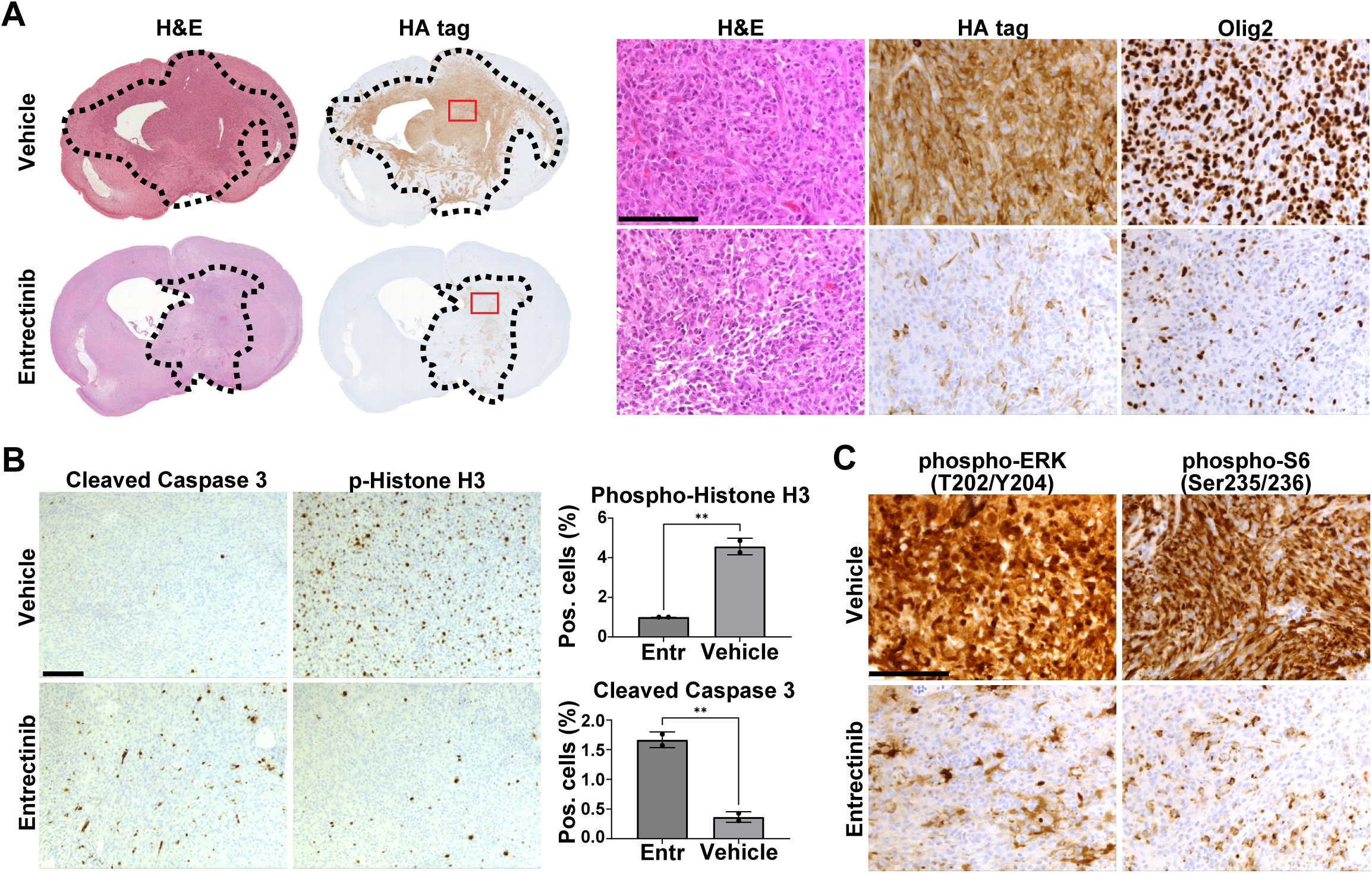
In vivo TKI treatment leads to the regression of *NTRK* gene fusion-driven mouse gliomas. A) Representative H&E and IHC (HA-tag, Olig2) images of HA-EML4-NTRK3-driven mouse gliomas treated with either Vehicle or Entrectinib (60 mg/kg) for 24 hours. Black dotted line indicates tumor boundaries. B) Left: Representative images of IHC stainings for Cleaved Caspase 3 and phospho-Histone H3 of HA-EML4-NTRK3-driven mouse gliomas treated with either Vehicle or Entrectinib (60 mg/kg) for 24 hours. Right: Quantification of Cleaved Caspase 3 and phospho-Histone H3 stainings (n = 2 per condition). C) Representative IHC stainings for phospho-ERK and phospho-S6 of HA-EML4-NTRK3-driven mouse gliomas treated with either Vehicle or Entrectinib (60 mg/kg) for 24 hours. Scale bar indicates 100 µm. Error bars show SD. Analysis was done using two-tailed t-test (B). (**) P < 0.01.

Taken together, our results suggest that TKI treatment can induce the regression of *NTRK* fusion-driven mouse gliomas in vivo within hours.

### TKI treatment significantly prolongs the survival of *NTRK* gene fusion-driven mouse gliomas, but tumors ultimately recur after treatment discontinuation

To assess if the two first-generation TKIs Larotrectinib and Entrectinib can prolong the survival of *NTRK* fusion-driven mouse gliomas in vivo and if there is a difference in the response between tumors induced by different *NTRK* fusions, we selected six *NTRK* fusions and intracranially induced gliomas by RCAS viral injection into either neonatal (*ETV6-N3*, *EML4-N3*) or adult (*TN1*, *CN1*, *GN2*, *NN2*) N/tv-a *Cdkn2a* null mice and performed preclinical in vivo trials. To assess the initial tumor size before treatment and ensure a distribution of similarly sized tumors into each treatment group, a first T2w MRI was performed 15-25 days post tumor induction (depending on the *NTRK* fusion, see Suppl. Methods) and mice were then treated with either Vehicle (DMSO), Larotrectinib (100 mg/kg), or Entrectinib (30 mg/kg) for 14 days (for dosing and toxicity experiments see Suppl. Methods). The day after the last treatment, the mice received a second T2w MRI and were subsequently observed for the onset of tumor-related symptoms. No MRIs were performed for *EML4-N3* mice, since these were too young at treatment start due to the aggressive growth of these tumors.

Treatment with either Larotrectinib or Entrectinib significantly extended survival compared to Vehicle treatment, however, treated tumors ultimately recurred upon treatment discontinuation in all mice due to the presence of residual disease and treatment-resistant persister cells (Figure 5A, Suppl. Figure S4A). The extent to which survival was prolonged differed between the different *NTRK* fusions and the two TKIs (Suppl. Figure S4B). The largest increase in survival was observed for *EML4-N3*-driven tumors treated with Larotrectinib (36 days versus 16 days (Vehicle) median survival, p < 0.0001) and *ETV6-N3*-driven tumors (57 days (Larotrectinib) and 53 days (Entrectinib) versus 41.5 days (Vehicle) median survival, p < 0.0001 and p = 0.0001 respectively). We observed that for some fusion-driven tumor types, a specific TKI was more effective. In *CN1*-driven tumors Entrectinib treatment led to a significant survival benefit over Larotrectinib treatment (48 versus 41 days median survival, respectively, p = 0.0003), whereas in *NN2* (56 versus 50 days median survival, respectively, p = 0.001), *ETV6-N3* (57 versus 53 days median survival, respectively, p = 0.05), and *EML4-N3*-driven tumors (36 versus 19 days median survival, respectively, p < 0.0001) Larotrectinib treatment led to a significantly better outcome compared to Entrectinib treatment. There was no significant difference between both TKIs at the tested concentrations for *TN1* and *GN2* fusion-driven tumors.

**Fig 5:**
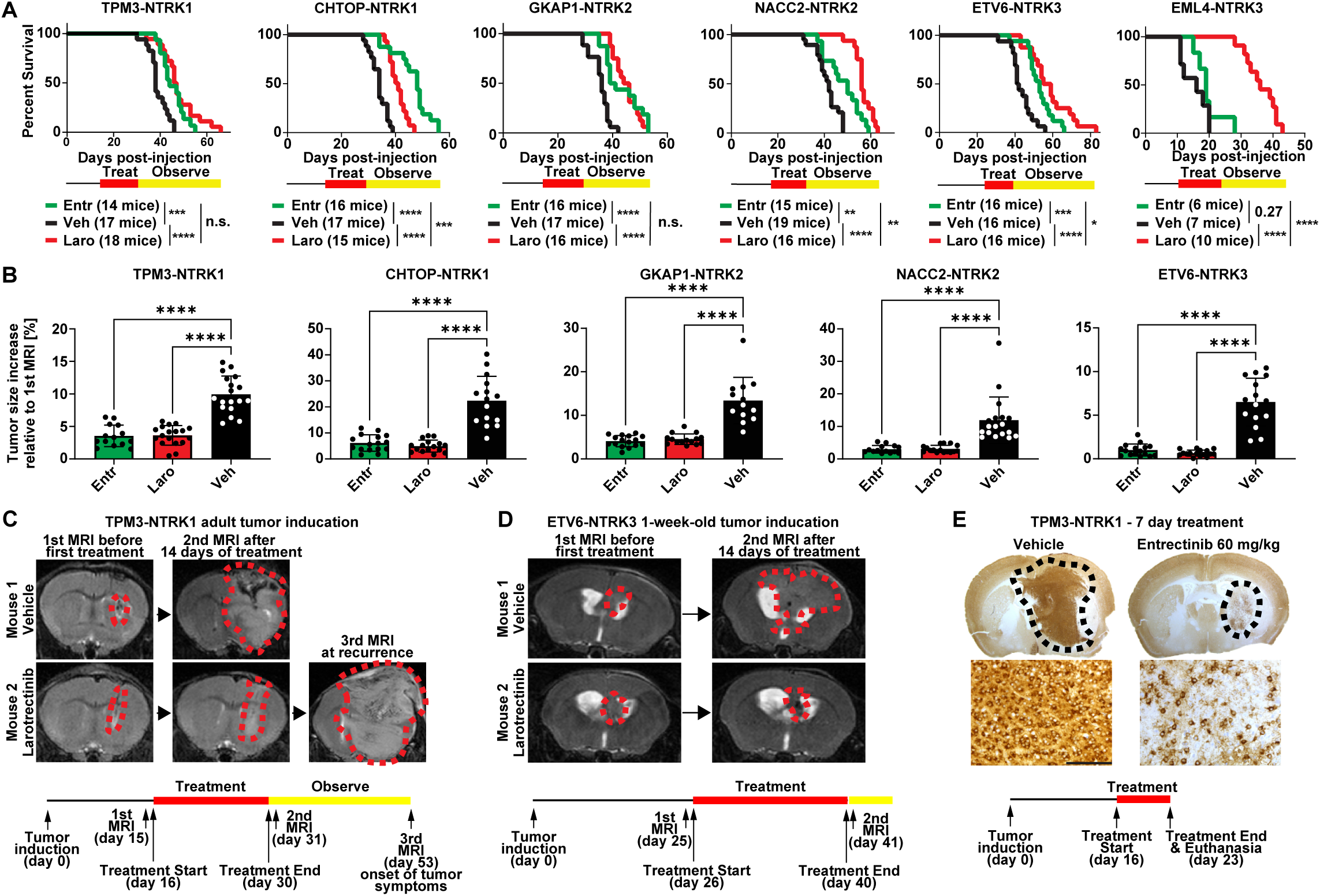
TKI treatment significantly prolongs the survival of NTRK gene fusion-driven mouse gliomas. A) Survival graphs of mice harboring *TPM3-NTRK1*, *CHTOP-NTRK1*, *GKAP1-NTRK2*, *NACC2-NTRK2*, *ETV6-NTRK3*, and *EML4-NTRK3*-driven gliomas treated with either Vehicle, Larotrectinib (100 mg/kg), or Entrectinib (30 mg/kg) via i.p. injection twice-a-day (b.i.d.) for 14 days. After treatment discontinuation, mice were observed for the onset of tumor-related symptoms. B) Graphs depicting change of tumor-size during the 14-day period of treatment with Vehicle, Larotrectinib, or Entrectinib. An initial MRI was performed one day before treatment start and a second MRI one day after treatment end. C-D) Representative MRIs of mice harboring *TPM3-NTRK1* (C)– or *ETV6-NTRK3* (D)-driven tumor treated with either Vehicle or Larotrectinib. E) Representative IHC stainings (TRK kinase domain) of *TPM3-NTRK1*-mouse gliomas after 7 days of treatment (i.p.; b.i.d.) with either Vehicle or Entrectinib (60 mg/kg). Red or black dotted line indicates tumor boundaries. Scale bar indicates 100 µm. Analysis was done using Log-rank (Mantel-Cox) test (A) or ordinary one-way ANOVA (B). (*) P < 0.05, (**) P < 0.01, (***) P < 0.001, (****) P < 0.0001.

We calculated the fold-change increase in tumor size based on pre– and post-treatment MRIs and observed that intra-parenchymal tumors were less susceptible to TKI treatment than extra-parenchymal tumors (Figure 5B). Extra-parenchymal (extra-axial and extracranial) tumors were almost uniformly present in Vehicle-treated mice, but frequently absent or smaller in TKI-treated mice (Suppl. Figure S4C-D). Intracranial tumors in Larotrectinib and Entrectinib-treated mice were significantly smaller compared to Vehicle-treated mice for each of the *NTRK1/2/3* fusion-driven glioma models at the end of treatment (Figure 5B-C). However, adult intracranial tumors (*TN1*, *CN1*, *GN2*, *NN2*) continued to grow while on treatment with Larotrectinib and Entrectinib and were between 3 to 6-fold larger compared to before treatment, albeit at a lower rate compared to Vehicle-treated tumors (10 to 22-fold larger compared to before treatment, depending on the tumor type). By contrast, pediatric *ETV6-N3* tumors remained stable or even shrank during TKI treatment; Larotrectinib and Entrectinib treatment resulted in a regression or stagnation of intra-parenchymal tumor growth (66% and 100% average tumor volume, respectively, relative to before treatment start), whereas Vehicle-treated tumors significantly grew (651% average tumor volume relative to the first MRI) (Figure 5D, Suppl. Figure S4E). Interestingly, the differences in survival between Larotrectinib and Entrectinib (such as for *CHTOP-NTRK1*) were not reflected by significantly different tumor sizes as measured by MRI.

As described above, we observed that similar to human *NTRK* fusion-positive tumors, TKI treatment led to tumor regression/stable disease or at least reduced tumor growth in our mouse models, but was unable to completely eradicate tumors, ultimately resulting in tumor recurrence once treatment was discontinued. Therefore, we sought to assess if an increased TKI dose would be sufficient to completely ablate all tumor cells in vivo. To this end, we treated *TN1*, *CN1*, and *GN2* tumor mice with 60 mg/kg Entrectinib b.i.d. (n = 2 per group) or Vehicle for 7 days. We had previously used a concentration of 30 mg/kg b.i.d. for the 14-day treatment since higher Entrectinib doses resulted in increased toxicity over longer treatment periods. While tumors in Entrectinib-treated mice were significantly smaller compared to Vehicle-treated tumors, treatment-resistance persister cells were present even at this increased dose, suggesting that TKI treatment alone is unable to completely eradicate these tumors in vivo (Figure 5E, Suppl. Figure S4F).

Taken together, our results show that treatment with Larotrectinib and Entrectinib can significantly prolong the survival of *NTRK* fusion-driven mouse gliomas in vivo, however, ultimately all tumors recurred upon treatment discontinuation due to the presence of persister cells. Tumors derived from different *NTRK* fusions do not respond equally to the two TKIs.

### Combination therapy of TRK and MEK inhibition is superior to TRK inhibition alone against NTRK fusion-driven cells

To investigate why treatment with Entrectinib was frequently less effective compared to Larotrectinib, we treated several mice harboring either ETV6-N3, TN1, or CN1 gliomas for 14 days with either Larotrectinib (100 mg/kg) or Entrectinib (30 mg/kg) for 14 days twice-a-day and euthanized them 1 hour after the last treatment. Treatment with either Larotrectinib or Entrectinib lead to a similar decrease of p-AKT levels in tumor tissues compared to Vehicle-treatment. However, p-ERK levels were considerably higher in tumor tissues of mice treated with Entrectinib compared to Larotrectinib, albeit lower than in Vehicle-treated mice (Figure 6A, Suppl. Figure S5A).

**Fig 6:**
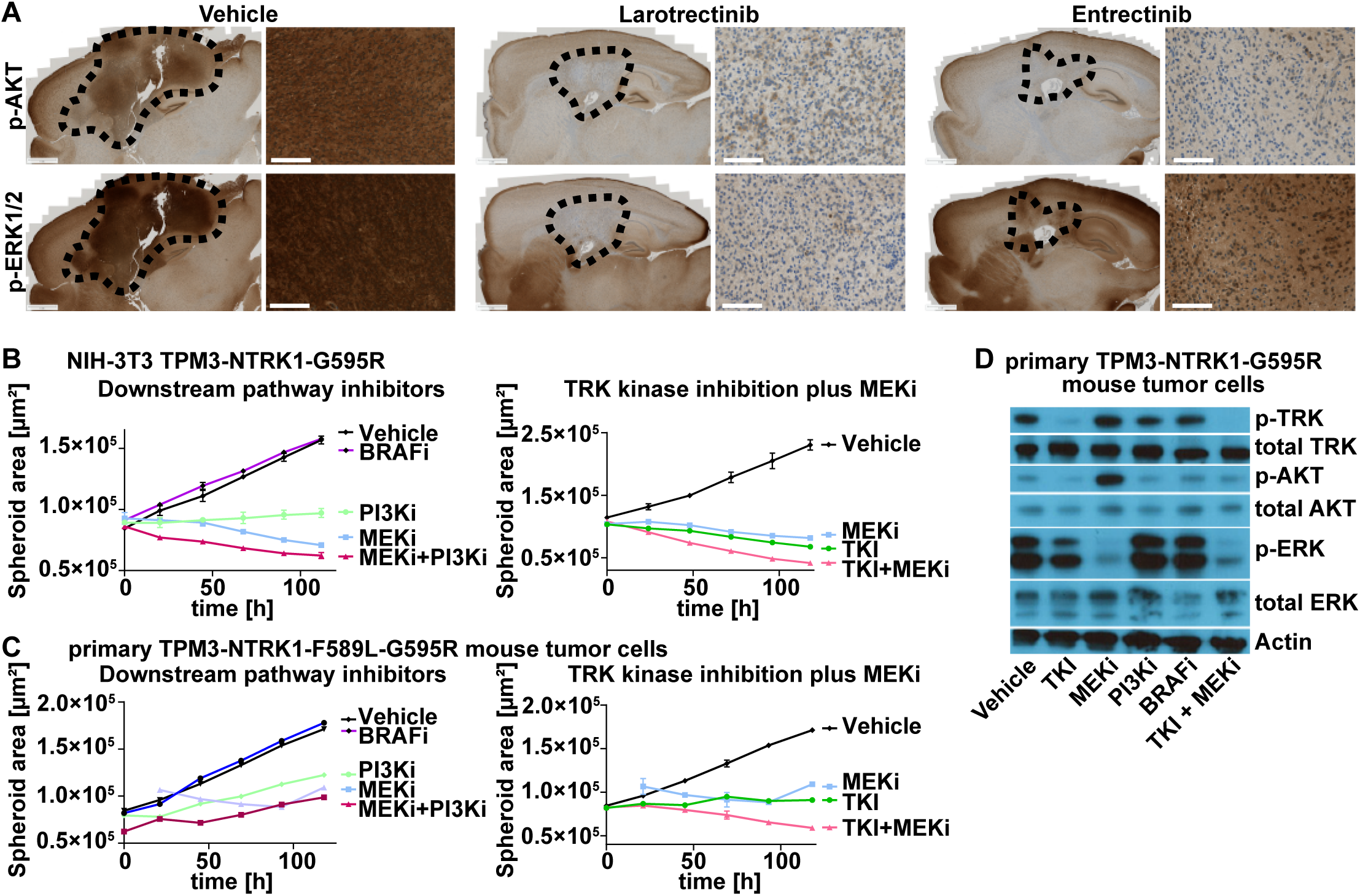
Combination therapy of TRK and MEK inhibition is superior to TRK inhibition alone against NTRK fusion-driven cells. A) Representative IHC stainings (phospho-AKT (Ser473); phospho-ERK1/2 (Thr202/Thr185)) of *ETV6-NTRK3*-mouse gliomas (Vehicle or 14 days of Larotrectinib (100 mg/kg) or Entrectinib (30 mg/kg) treatment). Black dotted line indicates tumor boundaries. Scale bars indicate 1 mm (whole brain slides) or 100 µm (high magnification panels). B-C) Growth curve of TPM3-NTRK1-G595R-expressing NIH-3T3-tv-a (C) or primary TPM3-NTRK1-F589L-G595R mouse tumor (D) cell spheroids treated with either DMSO (0.1%), Repotrectinib (TKI), Trametinib (MEKi), Taselisib (PI3Ki), PLX8394 (BRAFi), or a combination of Repotrectinib plus Trametinib (TKI + MEKi). All inhibitors were used at 100 nM except Trametinib at 10 nM. D) Representative Western Blot of primary TPM3-NTRK1-G595R mouse tumor cells treated with either DMSO (0.1%; Vehicle) or indicated inhibitors (same as C-D).

Since we observed increased p-ERK levels in tumors that tended to display a shorter survival, we speculated that ERK signaling plays a critical role in NTRK fusion-driven tumors. To test the efficacy of downstream pathway inhibition of both the PI3K-AKT-S6 and the RAF-MEK-ERK pathways we treated our previously established cell line models (in vitro-transduced NIH-3T3 cells and primary mouse tumor cells) with either Repotrectinib (TKI), Trametinib (MEKi), Taselisib (PI3Ki), or PLX3894 (BRAFi) and performed 3D spheroid assays. While BRAF inhibition had no effect, inhibition of both PI3K and MEK was able to inhibit the growth of NTRK fusion-driven cells (Figure 6B-C). Finally, combination treatment with TKIs and MEK inhibitors lead to a more pronounced growth inhibition than either alone (Figure 6B-C). TKI treatment decreased p-AKT levels but only partially reduced p-ERK levels, whereas MEK inhibition effectively reduced p-ERK levels but led to an increase in p-AKT levels. Combination treatment with TKIs and MEK inhibitors led to a decrease in both p-AKT and p-ERK levels (Figure 6D, Suppl. Figure S5B).

Taken together, these results suggest that ERK signaling plays an important role in NTRK fusion-driven tumors and might contribute to treatment resistance. In vitro, a combination of TKI and MEK inhibition is superior to either treatment alone.

## Discussion

Most adult IDH-wild type glioblastomas harbor a highly aberrant genome, leading to multiple concurrent and redundant drivers that make targeting of specific pathways difficult. By contrast, pediatric-type high-grade gliomas are enriched for gene fusions involving receptor tyrosine kinase genes (such as *NTRK1-3*, *ALK*, and *ROS1*) and harbor an otherwise relatively stable genome, making them ideal candidates for targeted therapy with tyrosine kinase inhibitors. These inhibitors show high initial response rates, frequently leading to either tumor regression or stable disease; however, surviving therapy-resistant persister cells eventually accumulate additional mutations, allowing them to regrow the tumor even in the presence of the kinase inhibitors ^3, 26^. The blood-brain barrier limits drug availability in the brain and brain tumors, further impeding the efficacy of these inhibitors.

Using the RCAS/tv-a system ^16^, we have developed a series of novel genetically engineered mouse models of pediatric-type *NTRK* fusion gene-driven high-grade gliomas. These GEMMs have several advantages over conventional xenograft or syngraft/allograft models and even over germline GEMMs. Our RCAS/tv-a tumor models are caused by somatic cell gene transfer of defined oncogenes (such as *NTRK* gene fusions) into *Nestin*-expressing neural stem and progenitor cells. Since these tumors arise de novo from within the mouse’s own cells, there is no potential graft versus host effect that might occur in allograft models, and in contrast to xenograft transplantation models these tumors can be induced in mice that harbor a completely intact immune system. An advantage over germline GEMMs is the flexibility of this system, that allows us to rapidly introduce additional mutations into the system to model for example treatment-experienced tumors, such as resistance-associated point mutations in the fusion protein kinase domain and test different *NTRK* fusion types. These models will help us to better understand the biology of treatment response and resistance to therapy of *NTRK* fusion-driven pediatric-type gliomas in a genetically-defined system with tumors that are caused and driven by the actual *NTRK* gene fusion, and that therefore directly rely on its activity.

All tested NTRK fusions were highly oncogenic when expressed in the brain, and in the absence of *Cdkn2a* expression resulted in uniform tumors that are ideal model systems for pre-clinical in vivo trials. Our data suggests that NTRK gene fusions indeed could be the initiating events in humans and is consistent with the relative lack of additional genetic rearrangement found in these tumors relative to adult gliomas. We also found that tumors were more aggressive when induced in younger mice (e.g. shortly after birth) compared to older mice, which may give an explanation why these gene fusions are enriched in tumors in infants and younger children.

There is a plethora of different *NTRK* fusions and clinical trials (either with *NTRK* fusion-positive brain or peripheral tumors) show that not all tumors respond to TKI treatment equally, sometimes even if they harbor the same gene fusion ^3, 8, 9^. This might be in part due to the presence of additional oncogenic driver mutations or the presence of kinase domain point mutations, but it is also unclear if different *NTRK* fusions (and the tumors that harbor these fusions) are equally susceptible to TKI treatment. All of these individual fusions are rare in humans, making trials comparing drugs against tumors with specific fusions impractical. Therefore, mouse models with a defined genetic background of tumors driven by specific gene fusions and their resistant mutations are needed.

Tumors induced by different NTRK fusions varied significantly in aggressiveness. Tumors induced by *EML4-NTRK3*, *BTBD1-NTRK3*, *CHTOP-NTRK1*, *TPM3-NTRK1*, and *GKAP1-NTRK2* were generally the most aggressive, whereas tumors induced by *ETV6-NTRK3* displayed the longest latency. This is especially noteworthy, since 1) *ETV6-NTRK3*, *EML4-NTRK3* and *BTBD1-NTRK3* all harbor the same NTRK3 kinase domain, and 2) *ETV6-NTRK3* is one of the most common *NTRK* fusions in pediatric-type gliomas and other peripheral tumors ^4, 5, 8^. It remains unclear why *ETV6-NTRK3* is less oncogenic compared to the other fusions and it might be speculated that the N-terminal fusion partner can influence the oncogenicity of the *NTRK* fusion, potentially through varying abilities to induce fusion dimerization necessary for autophosphorylation and activation ^27^.

All tested *NTRK* fusion-driven mouse gliomas responded to treatment with the two first-generation TKIs Larotrectinib and Entrectinib. We observed treatment responses within the first 24 hours of treatment, accompanied by the down-regulation of oncogenic PI3K-AKT-S6 and RAF-MEK-ERK signaling pathways in a majority of cells. Similar to *NTRK* fusion-positive tumors in patients, tumor clearing was incomplete and surviving persister cells led to the regrowth of tumors after treatment discontinuation. The adult-induced tumors even continued to grow while on treatment, albeit at a slower rate than Vehicle-treated tumors. Limited drug penetration into the brain and increased toxicity at higher concentrations of the two first-generation inhibitors likely contributed to the incomplete responses of intra-parenchymal brain tumors, since extra-cranial and extra-axial tumors showed a more complete response, although these tumors also regrew in some cases, suggesting that there were persister cells in the extra-cranial/-axial tumors as well. Future characterization of these treatment-resistant persister cells will be necessary and important in order to develop effective combination therapies to improve the efficacy of TKIs.

We observed reproducible differences in the therapeutic responses to TKIs between the various *NTRK* gene fusion-driven tumors at the tested inhibitor concentrations. *CHTOP-NTRK1*-driven tumors responded significantly better to Entrectinib over Larotrectinib treatment, while in *NACC2-NTRK2*, *ETV6-NTRK3*, and *EML4-NTRK3*-driven tumors Larotrectinib treatment led to a significantly longer survival compared to treatment with Entrectinib. It is unclear whether this effect is related to fusion or tumor biology. In our in vitro systems we observed that Entrectinib was more potent against *TPM3-NTRK1*-expressing cells compared to *GKAP1-NTRK2* and *ETV6-NTRK3*-expressing cells. In general, Larotrectinib (100 mg/kg) was better tolerated than Entrectinib (30 mg/kg) by tumor-bearing mice, limiting the active concentration that could be administered. We observed increased p-ERK levels in tumors of Entrectinib-treated mice, suggesting that ERK signaling might contribute to resistance to TKI therapy. In vitro, MEK inhibition with Trametinib was able to inhibit the growth of NTRK fusion-driven cells and a combination of TKIs and MEK inhibitors was superior to either treatment alone.

In summary, this study highlights the utility of genetically engineered mouse models of specific oncogenic drivers to optimize therapy of tumors driven by specific molecular mechanisms where clinical trials are not possible because of their rarity. These models will be valuable tools for the study of treatment response and the development of resistance to therapy in these tumors.

## Materials & Methods

### Generation of RCAS mouse tumors

All animal experiments were done in accordance with protocols approved by the Institutional Animal Care and Use Committees of Fred Hutchinson Cancer Center (protocol no. 50842) and followed National Institutes of Health guidelines for animal welfare. The RCAS/tv-a system used in this work has been described previously ^20^. *Nestin* (N)/tv-a;*Cdkn2a*-wild type and null mice were used for RCAS-mediated brain tumor formation in this study and have been described previously ^20^. DF1 cells (1 × 10^5^) in a volume of 1 μL were injected near the ventricles into newborn (p0-p2) or one-week-old (p7) pup brains or into the striatum of 5-7-week-old adult mice as described previously ^20, 28^. For peripheral injections, DF1 cells (1 × 10^5^) in a volume of 1-2 μL were injected intraperitoneally (i.p.). Mice were monitored until they developed symptoms of disease, such as visible tumors, lethargy, poor grooming, weight loss, dehydration, macrocephaly, seizures, jumping, or paralysis, or until a predetermined study end point.

### In vivo TKI treatment

For in vivo treatment, inhibitors were resuspended in DMSO and diluted to a final buffer solution of 10% DMSO, 10% EtOH, 30% Solutol (Sigma Aldrich 42966), and 50% Tartaric acid buffer. Tartaric acid buffer stock was prepared by mixing 10 ml of 0.15 M DL-tartaric acid (Sigma Aldrich PHR1472) with 50 ml of 0.15 M tartrate sodium (Sigma Aldrich PHR1409). 10 ml of the Tartaric acid buffer stock were subsequently mixed with 40 ml of Saline Solution (0.9%) to create the final Tartaric acid buffer. Mice were treated with either Vehicle (10% DMSO), Larotrectinib (100 mg/kg), or Entrectinib (30 or 60 mg/kg) at 10 ul per gram i.p. injection twice-a-day (b.i.d.) for 14 days.

See also supplemental experimental procedures.

## Funding

National Institutes of Health grant U54 CA243125 (ECH)

National Institutes of Health grant R35 CA253119-01A1 (ECH)

Preclinical Imaging Shared Resource RRID:SCR_022616

National Institutes of Health grant P30 CA015704 (Fred Hutch/University of Washington Cancer Consortium).

National Institutes of Health Shared Instrumentation Grant S10OD26919 (Fred Hutch/University of Washington Cancer Consortium).

This project has been funded in whole or in part with Federal funds from the National Cancer Institute, National Institutes of Health, under Contract No. 75N91019D00024, Task Order No Q3. The content of this publication does not necessarily reflect the views or policies of the Department of Health and Human Services, nor does mention of trade names, commercial products or organizations imply endorsement by the U.S. Government.

## Author Contributions

Conceptualization SS, ZRR, DKK, REP, ECH, FS. Performed experiments SS, ZRR, ASY, MW, JW, DR, JZY, FS. Data analysis SS, ZRR, ASY, AGP, DCQ, BNG, NA, PJC, FS. Original manuscript writing SS, PJC, ECH, FS. Review and editing ECH, FS. Funding acquisition ECH. Supervision REP, ECH, FS. All authors read, reviewed, and approved the manuscript.

## Acknowledgements

We thank Denis Adair, Linda Lew, and Kelly Grissom for continued administrative assistance and support throughout these experiments. We thank Elizabeth Jensen and Dolores Covarrubias at the Fred Hutchinson Genomics Core for help with DNA sequencing. We thank Brianna Wrightson and Elena Carlson for developing and performing the MRI scans on tumor-bearing mice.

## Data and materials availability

The data that support the findings of this study are included with the manuscript and supplemental data files and are also available from the corresponding author upon reasonable request.

## Supplementary Information

Supplementary Figures S1-S5

Supplementary Figure Legends

Supplementary Methods

Supplementary Table S1

