## Supplementary material for "ERK signaling promotes resistance to TRK kinase inhibition in NTRK fusion-driven glioma mouse models": Suppl. Data

Supplementary Information

Table of Contents:

Supplementary Figures S1-S5
Supplementary Figure Legends
Supplementary Methods
Supplementary References

#### Suppl. Figure S1A-C

**A**

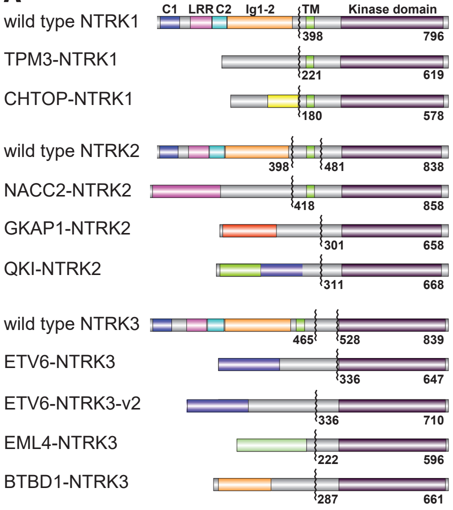


**B**

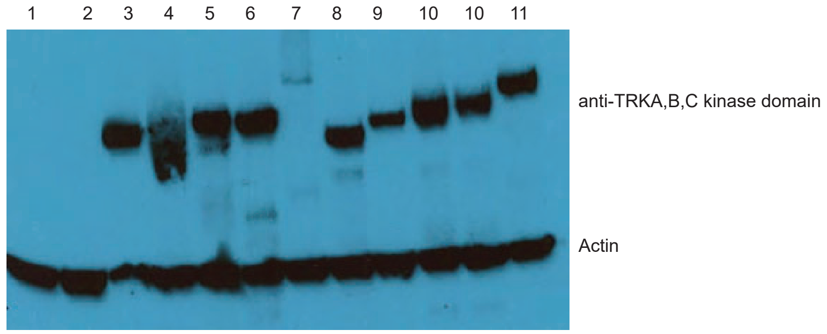


1 - untransfected
2 - RCAS-GFP
3 - RCAS-TPM3-NTRK1
4 - RCAS-CHTOP-NTRK1
5 - RCAS-GKAP1-NTRK2
6 - RCAS-QKI-NTRK2
7 - RCAS-NACC2-NTRK2
8 - RCAS-EML4-NTRK3
9 - RCAS-BTBD1-NTRK3
10 - RCAS-ETV6-NTRK3
11 - RCAS-ETV6-NTRK3-v2

**C**

| Nestin/tv-a wild type intracranial injection | | | | | | |
|---|---|---|---|---|---|---|
| Injection age | Newborn pups (p0-p2) | | 1-week-old pups (p7) | | Adult mice (5–7-week-old) | |
| Gene fusion | Tumor Penetrance | Median symptom-free survival | Tumor Penetrance | Median symptom-free survival | Tumor Penetrance | Median symptom-free survival |
| TPM3-NTRK1 | 6/6 mice (100 percent) | 16 days (hydrocephalus) | 23/27 mice (85 percent) | Mostly non-symptomatic 120 days post-inj. | 0/3 mice (0 percent) | Euthanized 120 days post-inj. |
| CHTOP-NTRK1 | 7/7 mice (100 percent) | 20 days | 5/5 mice (100 percent) | Mostly non-symptomatic 120 days post-inj. | 2/3 mice (67 percent) | Non-symptomatic 120 days post-inj. |
| GKAP1-NTRK2 | 5/5 mice (100 percent) | Mostly non-symptomatic 120 days post-inj. | 3/3 mice (100 percent) | Non-symptomatic 120 days post-inj. | 0/3 mice (0 percent) | Euthanized 120 days post-inj. |
| NACC2-NTRK2 | 6/7 mice (86 percent) | Non-symptomatic 120 days post-inj. | 4/4 mice (100 percent) | Non-symptomatic 120 days post-inj. | 0/1 mice (0 percent) | Euthanized 120 days post-inj. |
| QKI-NTRK2 | 5/5 mice (100 percent) | Non-symptomatic 120 days post-inj. | 1/5 mice (20 percent) | Non-symptomatic 120 days post-inj. | 0/2 mice (0 percent) | Euthanized 120 days post-inj. |
| ETV6-NTRK3 | 9/10 mice (90 percent) | Non-symptomatic 120 days post-inj. | 3/6 mice (50 percent) | Non-symptomatic 120 days post-inj. | n.d. | |
| EML4-NTRK3 | 4/4 mice (100 percent) | Non-symptomatic 120 days post-inj. | 4/4 mice (100 percent) | Mostly non-symptomatic 120 days post-inj. | n.d. | |
| BTBD1-NTRK3 | 6/7 mice (86 percent) | Non-symptomatic 120 days post-inj. | 6/6 mice (100 percent) | Non-symptomatic 120 days post-inj. | n.d. | |

### Suppl. Figure S1D

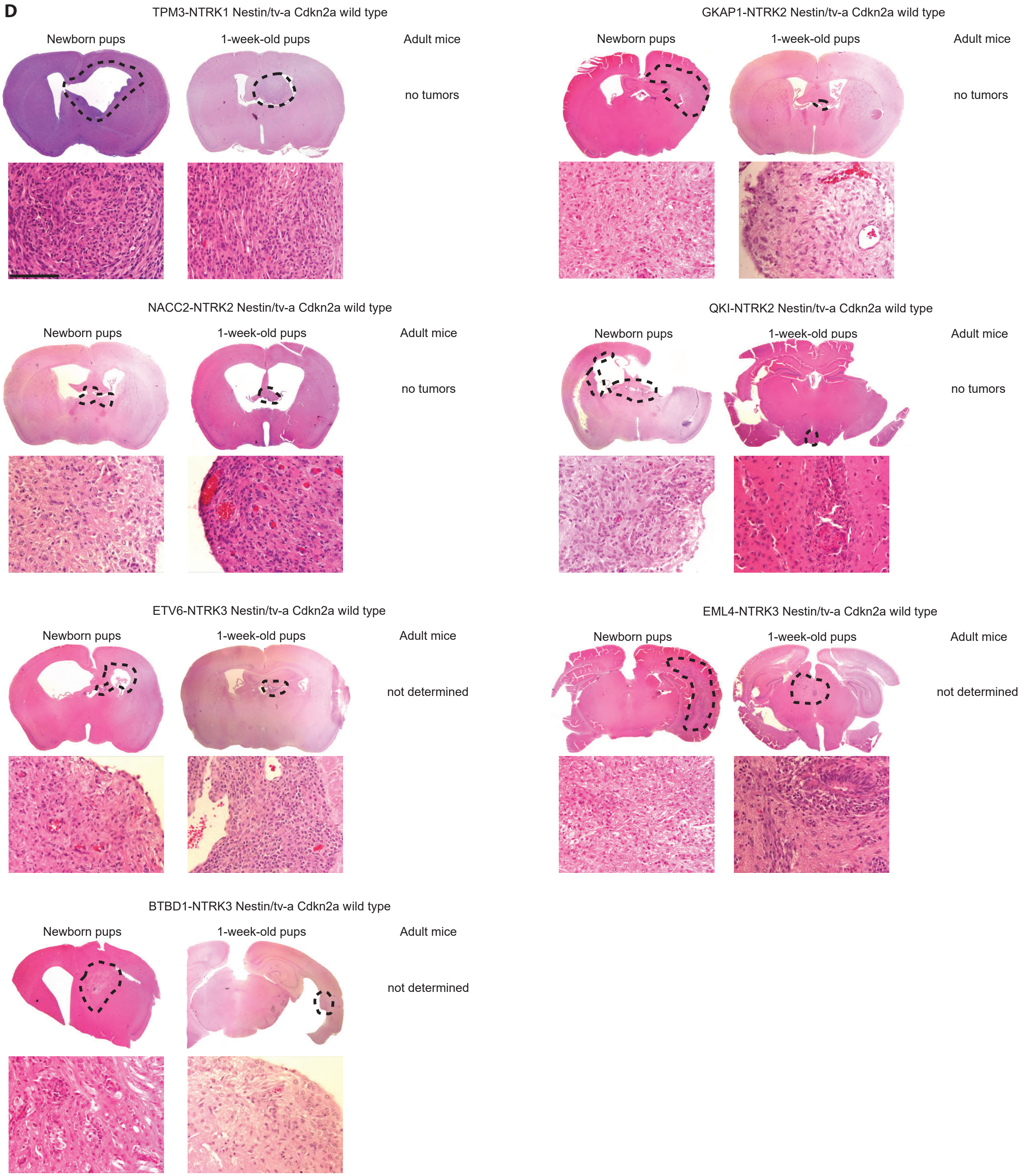

#### Suppl. Figure S1E-G

**E**

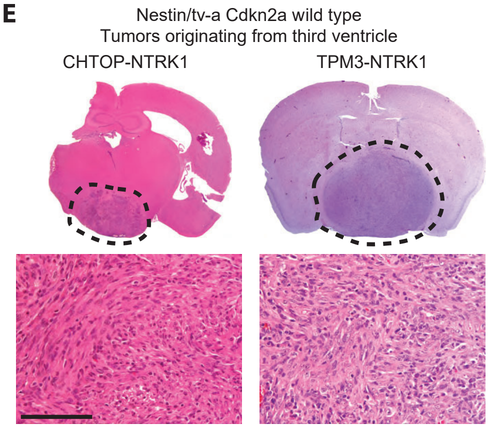


**F**

| Nestin/tv-a Cdkn2a null intracranial injection | | | | | | |
|---|---|---|---|---|---|---|
| Injection age | Newborn pups (p0-p2) | | 1-week-old pups (p7) | | Adult mice (5–7-week-old) | |
| Gene fusion | Tumor Penetrance | Median symptom-free survival | Tumor Penetrance | Median symptom-free survival | Tumor Penetrance | Median symptom-free survival |
| TPM3-NTRK1 | 4/4 mice (100 percent) | 14 days (hydrocephalus) | 6/6 mice (100 percent) | 21 days | 5/5 mice (100 percent) | 26 days |
| CHTOP-NTRK1 | 5/5 mice (100 percent) | 15 days (hydrocephalus) | 6/6 mice (100 percent) | 20 days | 5/5 mice (100 percent) | 27 days |
| GKAP1-NTRK2 | 8/8 mice (100 percent) | 20 days (hydrocephalus) | 10/10 mice (100 percent) | 21 days | 5/5 mice (100 percent) | 27 days |
| NACC2-NTRK2 | 8/8 mice (100 percent) | 17 days (hydrocephalus) | 8/8 mice (100 percent) | 24 days | 5/5 mice (100 percent) | 42 days |
| QKI-NTRK2 | 8/8 mice (100 percent) | 16 days (hydrocephalus) | 3/3 mice (100 percent) | 23 days | 5/5 mice (100 percent) | 41 days |
| ETV6-NTRK3 | 15/15 mice (100 percent) | 18 days (hydrocephalus) | 12/12 mice (100 percent) | 38 days | 5/5 mice (100 percent) | 53 days |
| EML4-NTRK3 | 6/6 mice (100 percent) | 16 days (hydrocephalus) | 7/7 mice (100 percent) | 17 days | 5/5 mice (100 percent) | 28 days |
| BTBD1-NTRK3 | 5/5 mice (100 percent) | 15 days (hydrocephalus) | 6/6 mice (100 percent) | 17 days | 5/5 mice (100 percent) | 28 days |

**G** Intracranial injection Nestin/tv-a Cdkn2a null adult mice

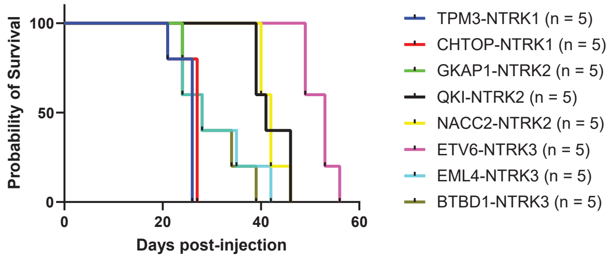

**Suppl. Figure S1H**

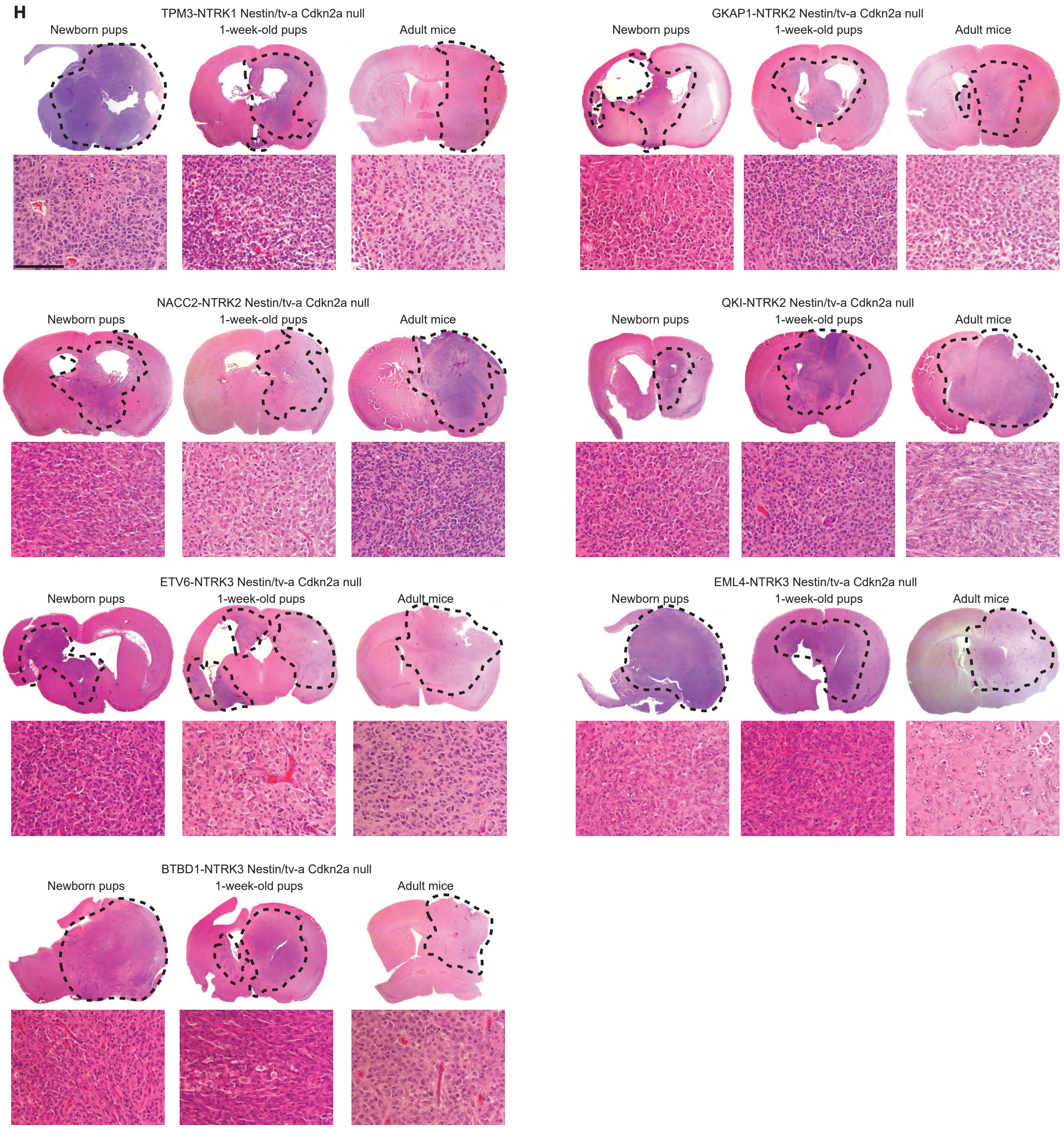

#### Suppl. Figure S1I-M

**I** Intracranial injection ETV6-NTRK3 variants
Nestin/tv-a Cdkn2a null 1-week-old mice

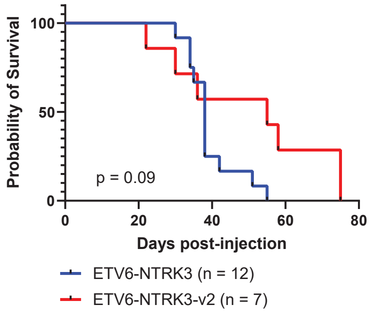


**J** Intracranial injection Nestin/tv-a Cdkn2a null newborn pups

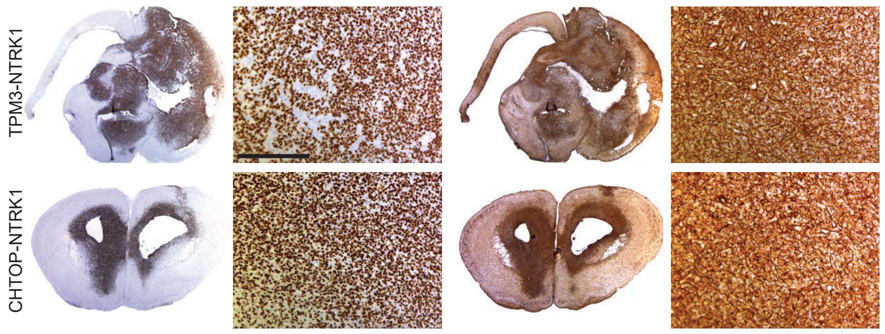


**K**

| Nestin/tv-a wild type + Pten knockdown intracranial inj. | | |
|---|---|---|
| Injection age | 1-week-old pups (p7) | |
| Gene fusion | Tumor Penetrance | Median symptom-free survival |
| TPM3-NTRK1 + shPten | 5/5 mice (100 percent) | Mostly non-symptomatic 120 days post-inj. |
| CHTOP-NTRK1 + shPten | 6/6 mice (100 percent) | 120 days |

**L**

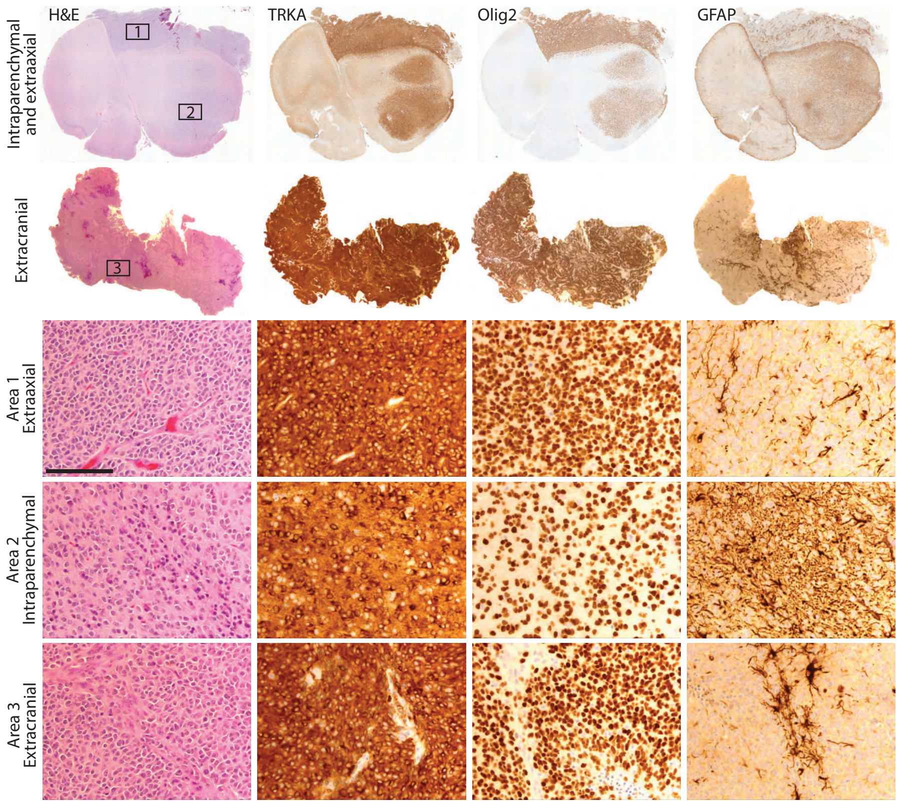


**M**

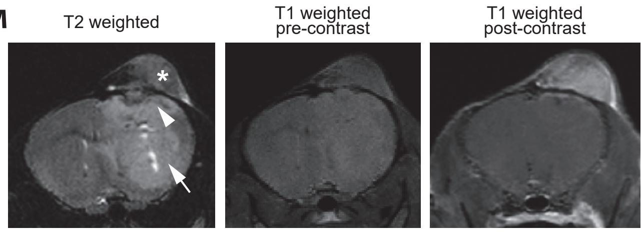

#### Suppl. Figure S1N-T

**N** Injection summary of intra-peritoneal RCAS injections

| Nestin/tv-a Cdkn2a null | | |
|---|---|---|
| Injection age | Newborn pups (p0-p2) | |
| Gene fusion | Tumor Penetrance | Median symptom-free survival |
| TPM3-NTRK1 | 11/11 mice (100 percent) | 30 days |

**O**

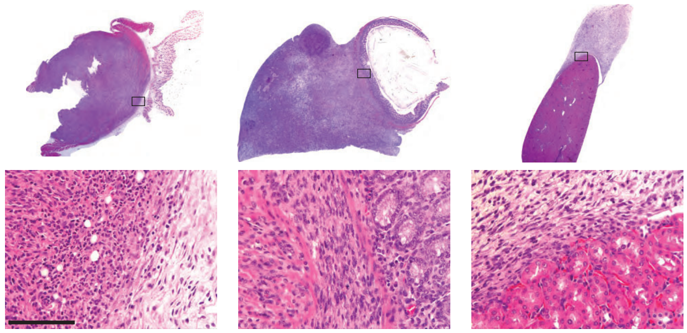


**P**

| Resistance-associated kinase point mutants (1-week-old pup intracranial injections (p7) | | | | |
|---|---|---|---|---|
| Background | Nestin/tv-a wild type | | Nestin/tv-a Cdkn2a null | |
| Gene fusion | Tumor Penetrance | Median symptom-free survival | Tumor Penetrance | Median symptom-free survival |
| TPM3-NTRK1 (unmutated) | 23/27 mice (85 percent) | Mostly non-symptomatic 120 days post-inj. | 6/6 mice (100 percent) | 21 days |
| TPM3-NTRK1-F589L | 6/10 mice (60 percent) | Non-symptomatic 120 days post-inj. | 5/5 mice (100 percent) | 19 days |
| TPM3-NTRK1-G595R | 4/6 mice (67 percent) | Mostly non-symptomatic 120 days post-inj. | 10/10 mice (100 percent) | 17 days |
| TPM3-NTRK1-F589L-G595R | 14/14 mice (100 percent) | Mostly non-symptomatic 120 days post-inj. | 14/14 mice (100 percent) | 20 days |
| TPM3-NTRK1-G667C | 6/6 mice (100 percent) | Mostly non-symptomatic 120 days post-inj. | 3/3 mice (100 percent) | 27 days |

**Q** Intracranial injection Nestin/tv-a Cdkn2a wild type One-week-old mice

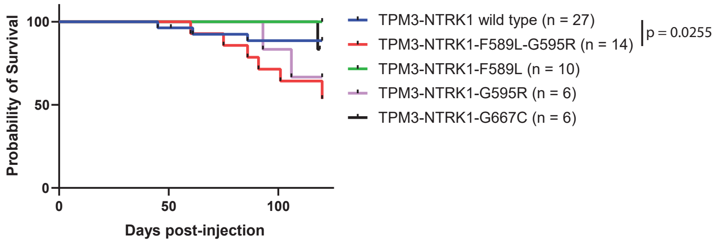


**R** Intracranial injection Nestin/tv-a Cdkn2a null One-week-old mice

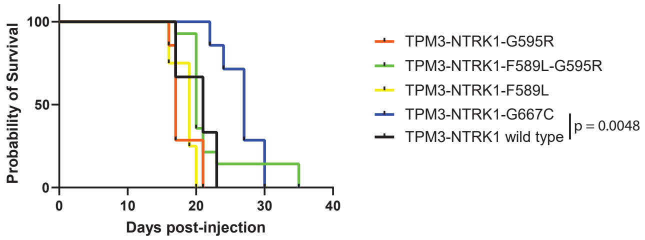


**S** Intracranial injection Nestin/tv-a Cdkn2a null One-week-old mice

**T** Intracranial injection Nestin/tv-a Cdkn2a wild type One-week-old mice

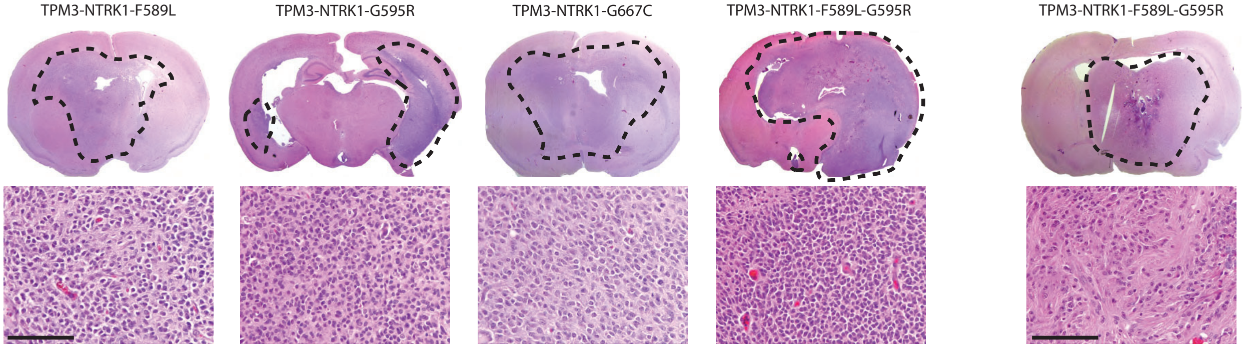

### Figure S2A TPM3-NTRK1

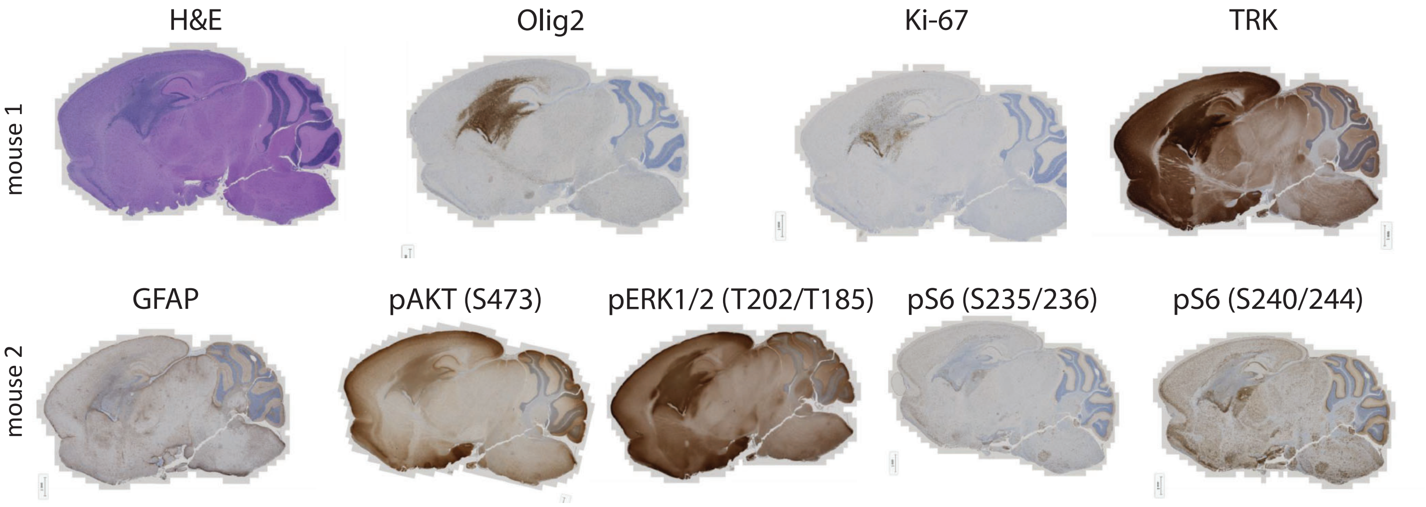

### Figure S2B CHTOP-NTRK1

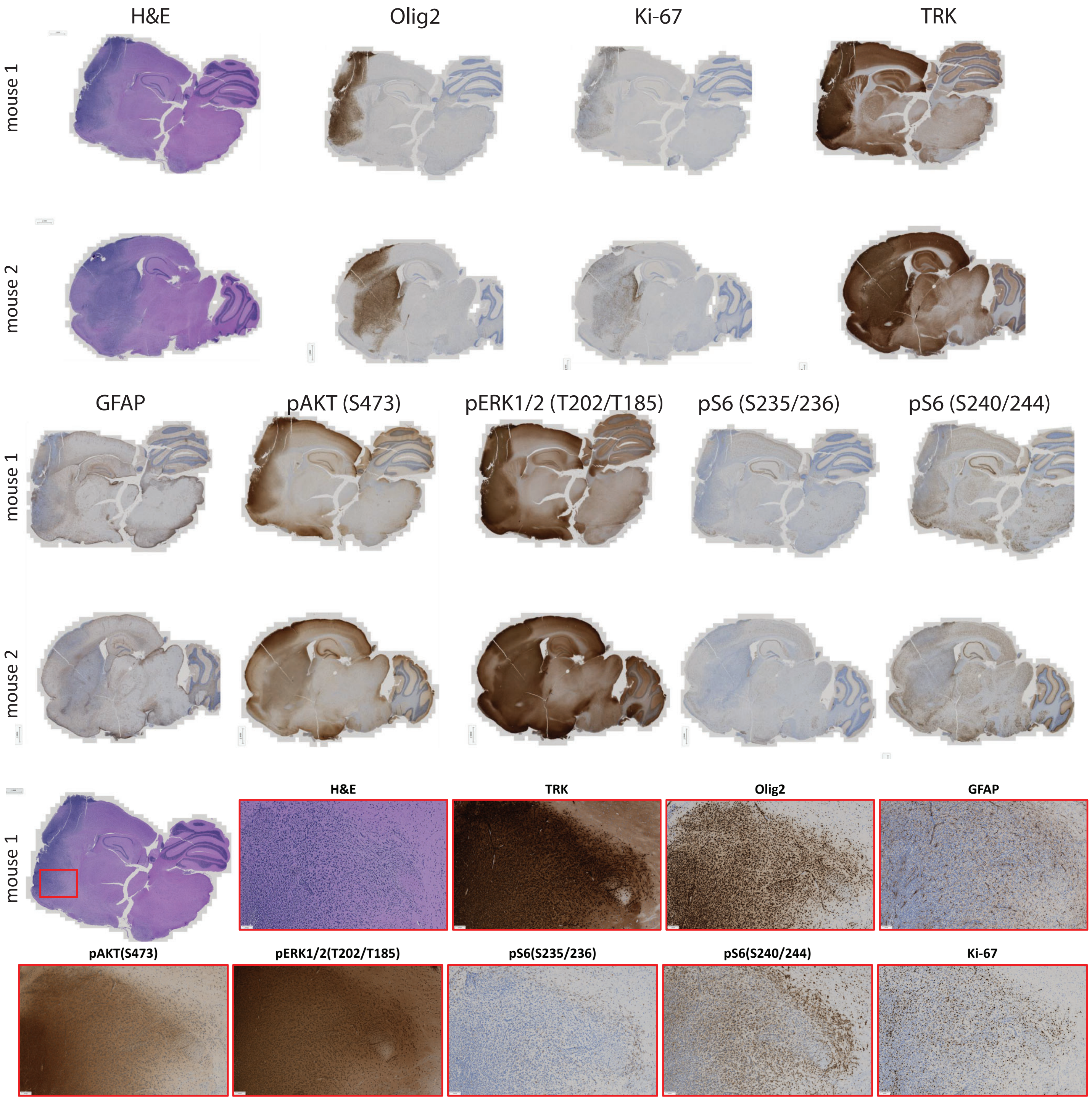

### Figure S2C GKAP-NTRK2

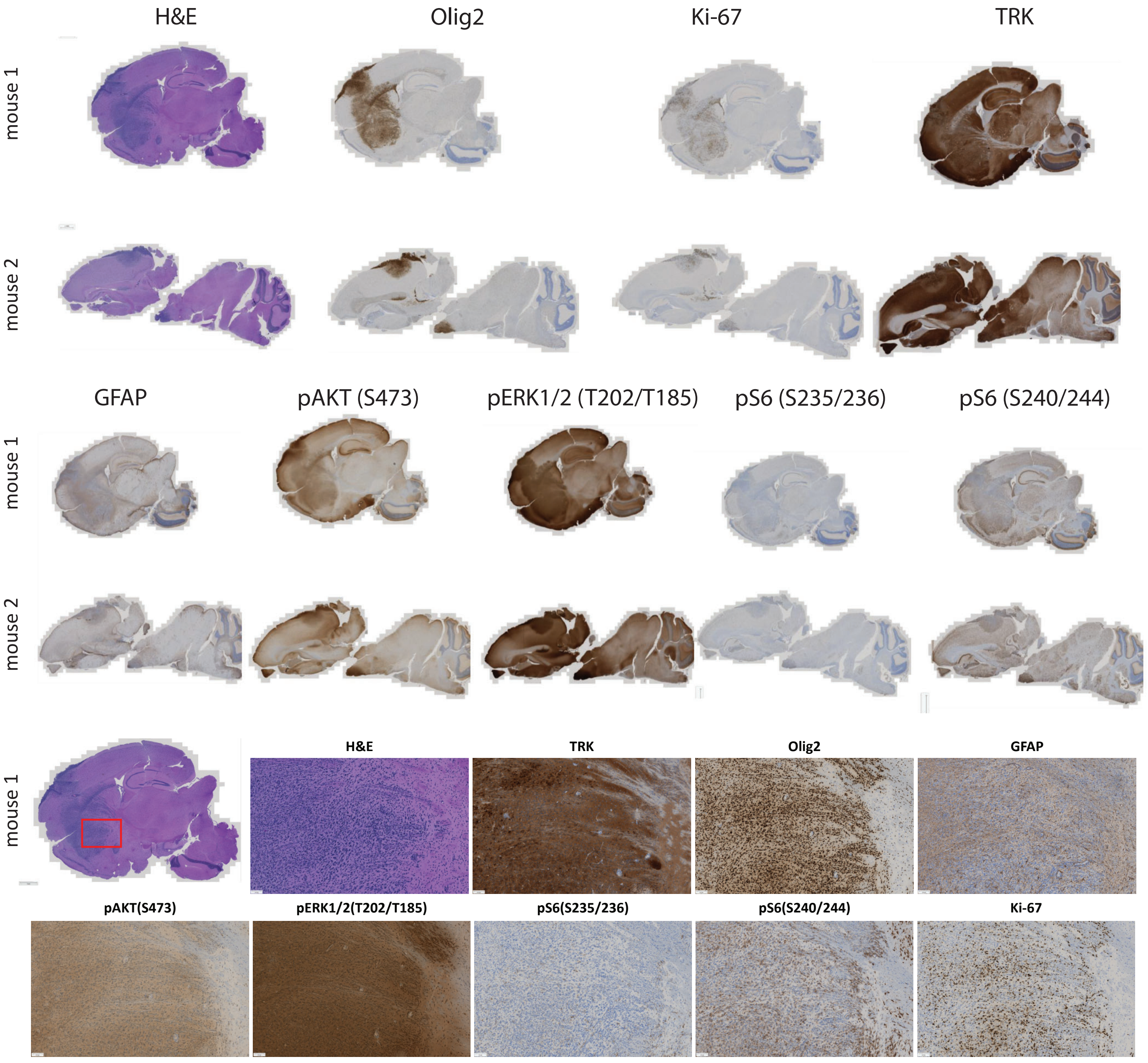

### Figure S2D QKI-NTRK2

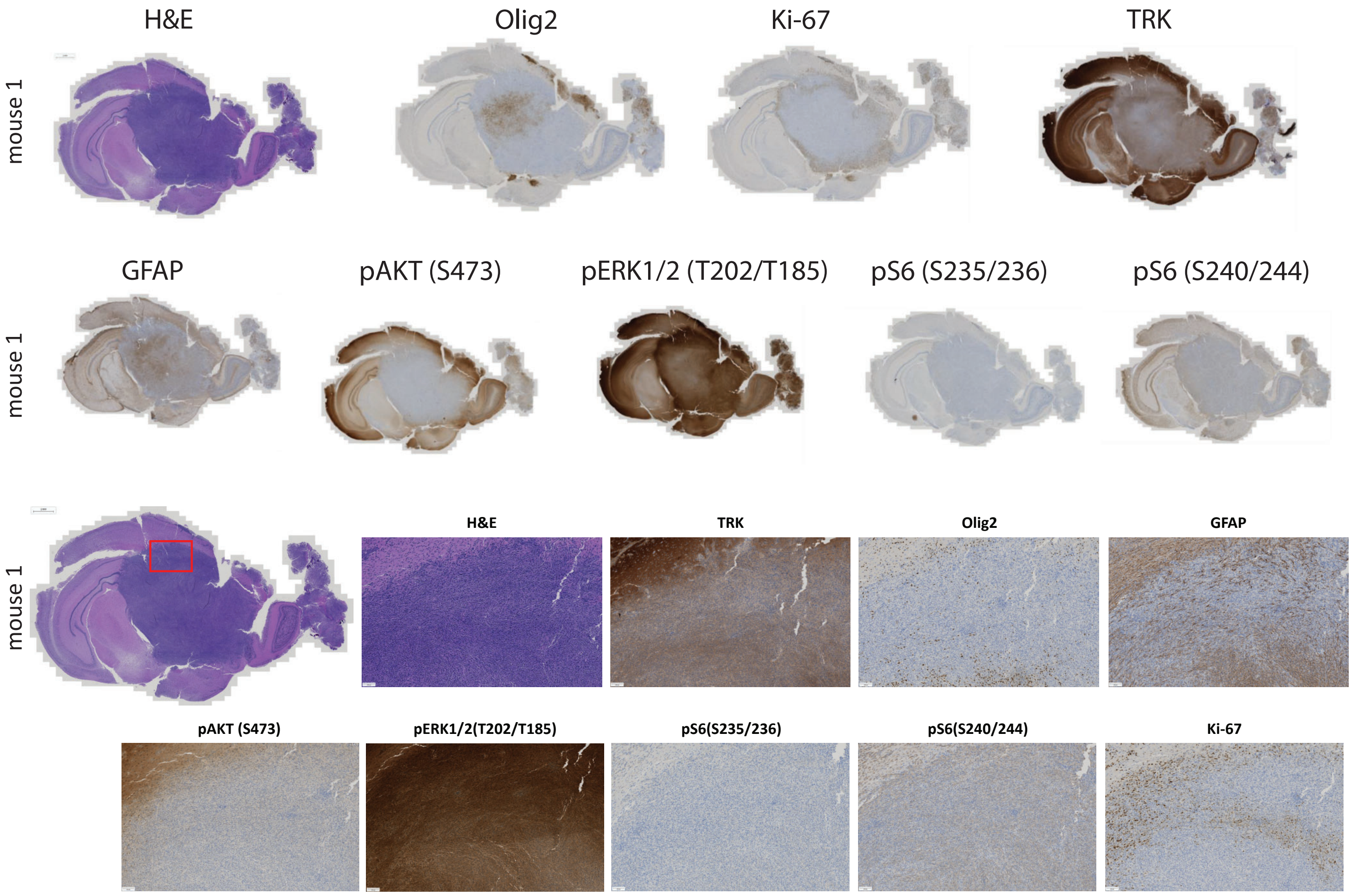

### Figure S2E NACC2-NTRK2

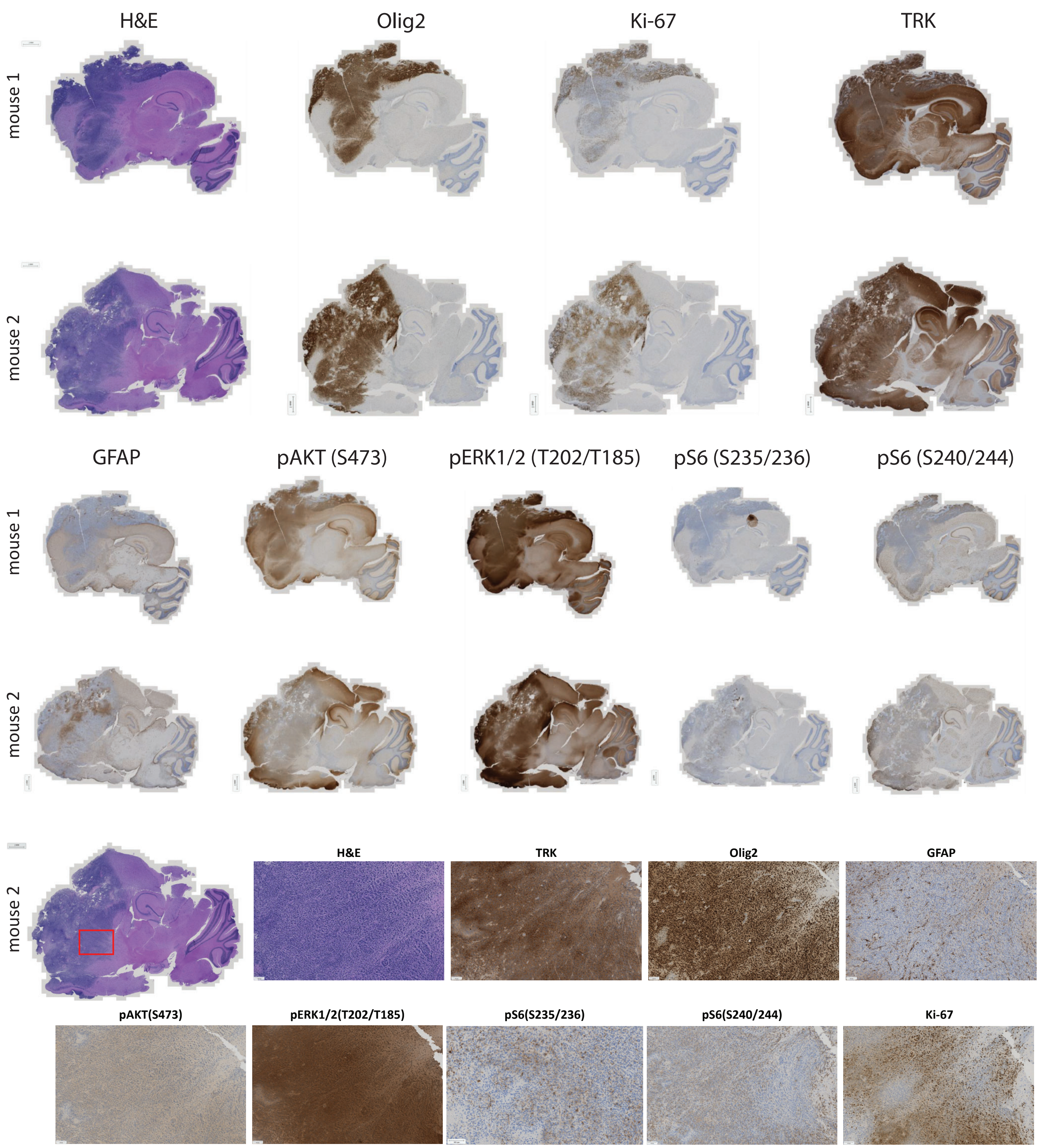

### Figure S2F ETV6-NTRK3

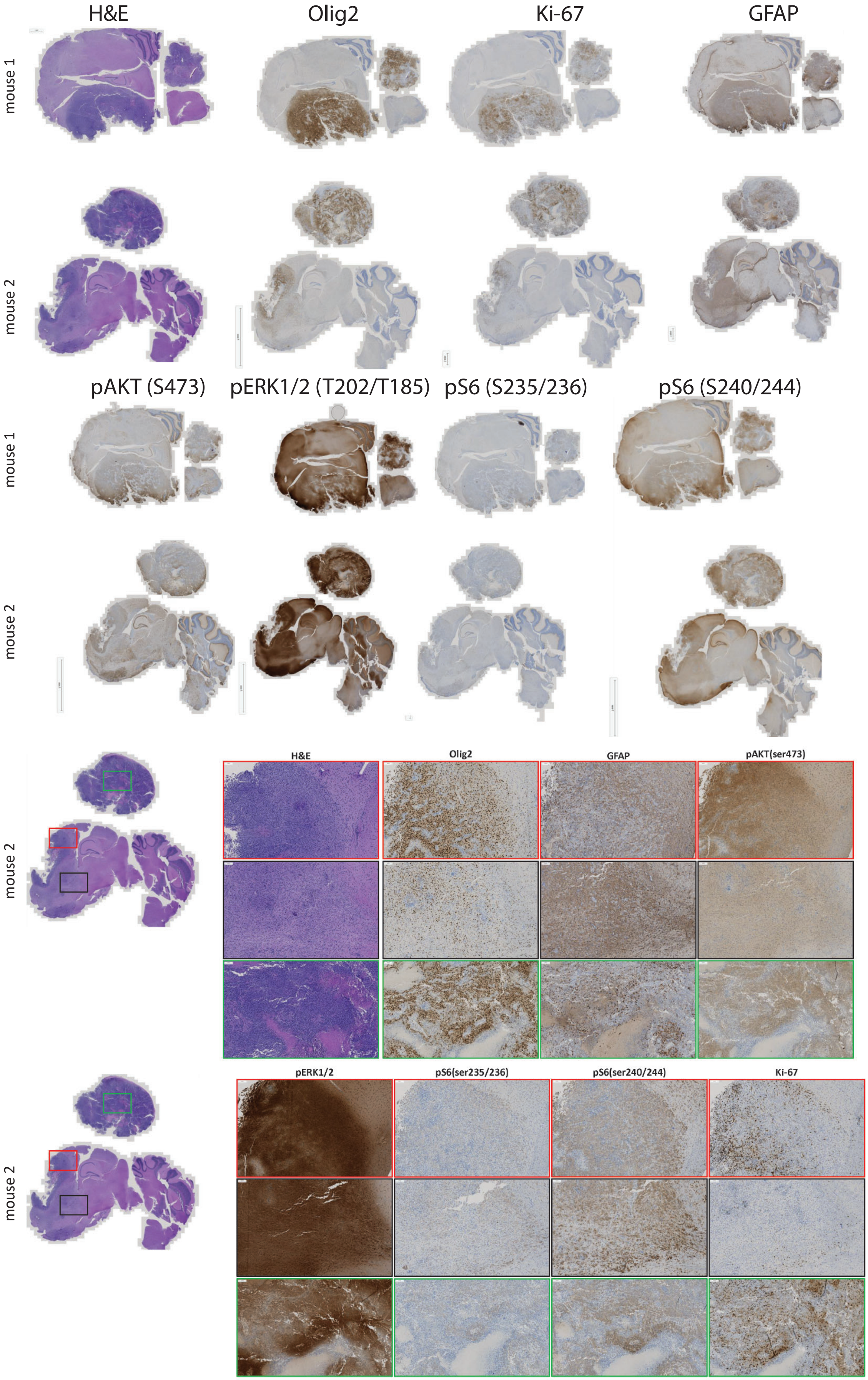

### Figure S2G BTBTD1-NTRK3

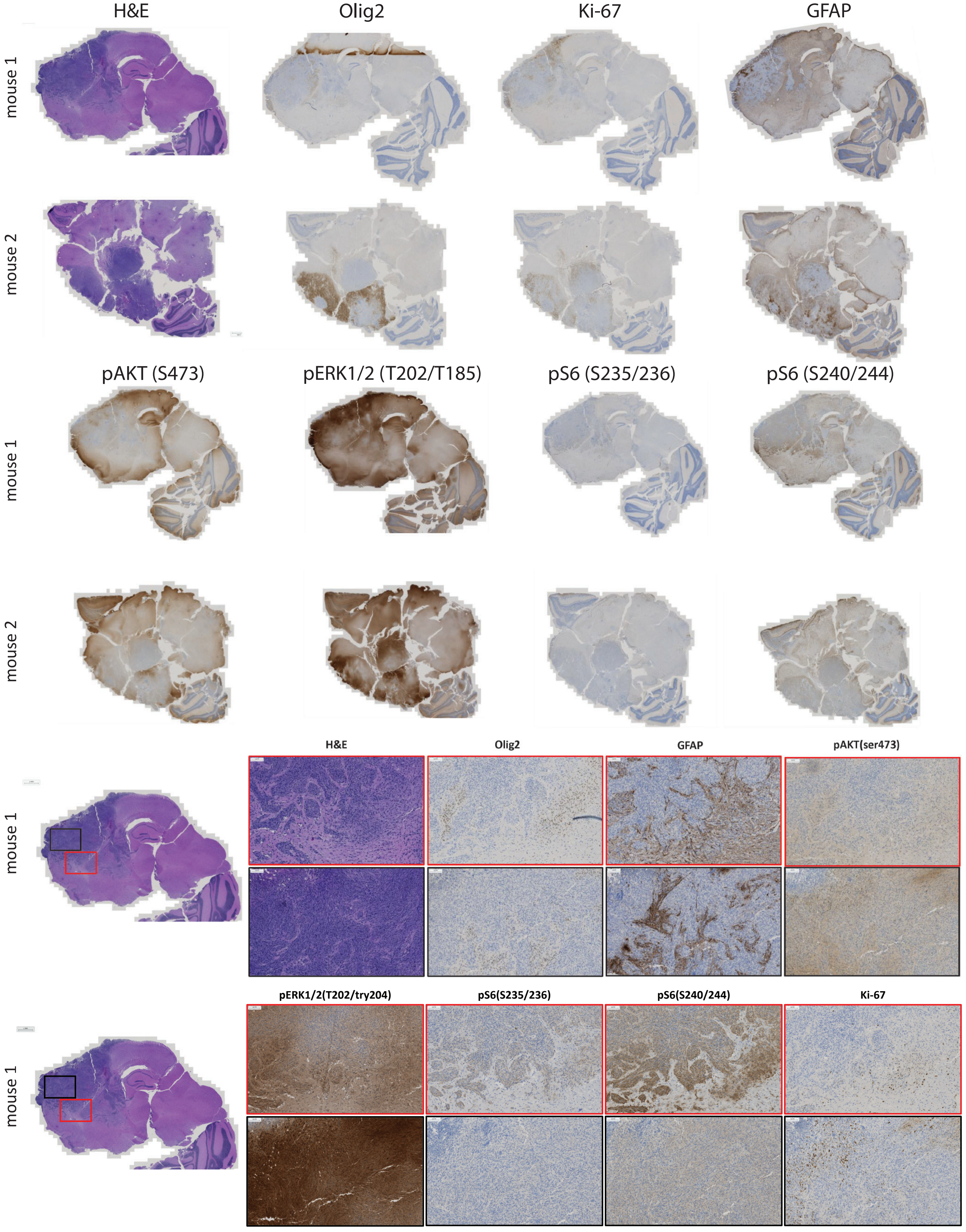

### Figure S2H EML4-NTRK3

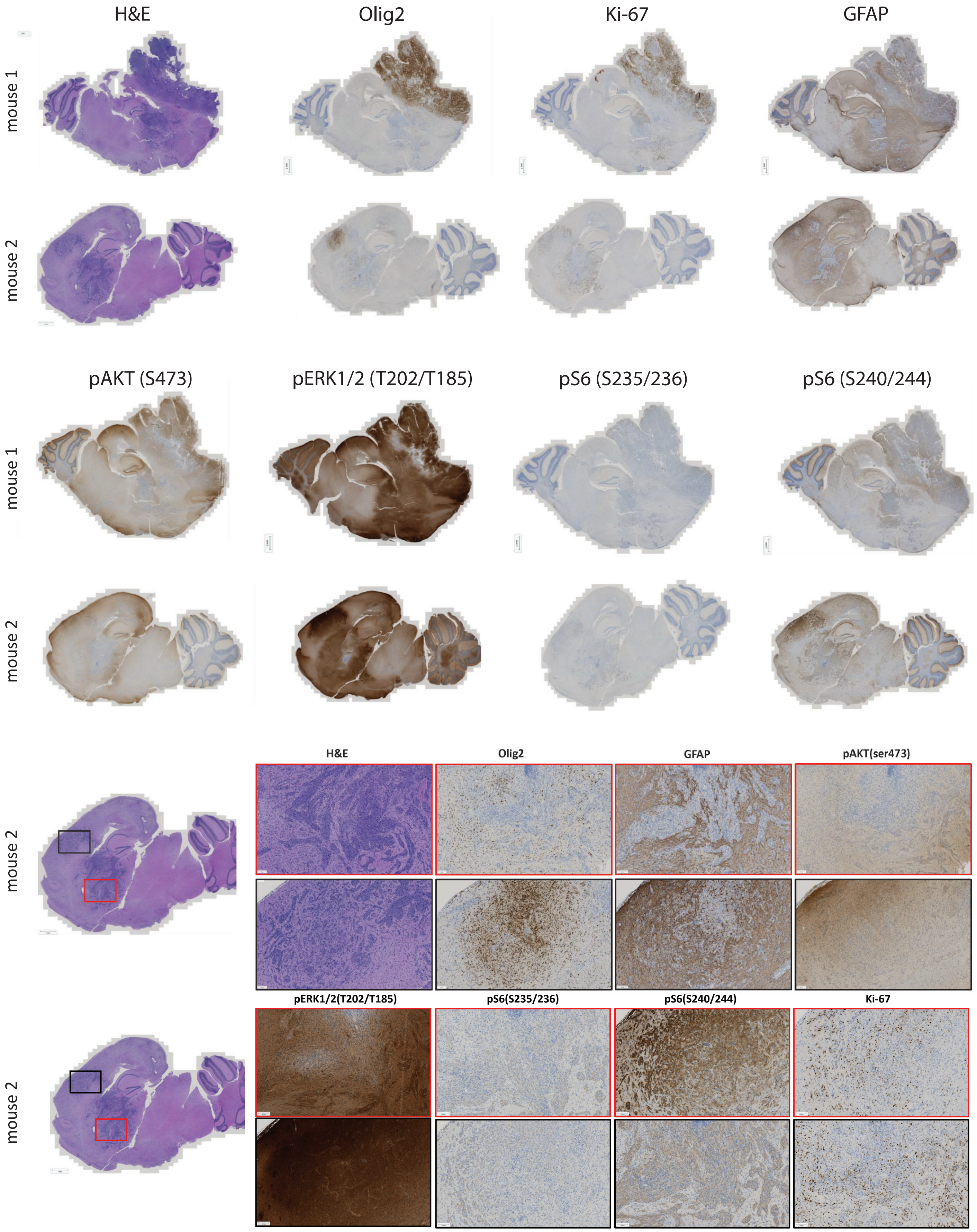

### Suppl. Figure S3A-D

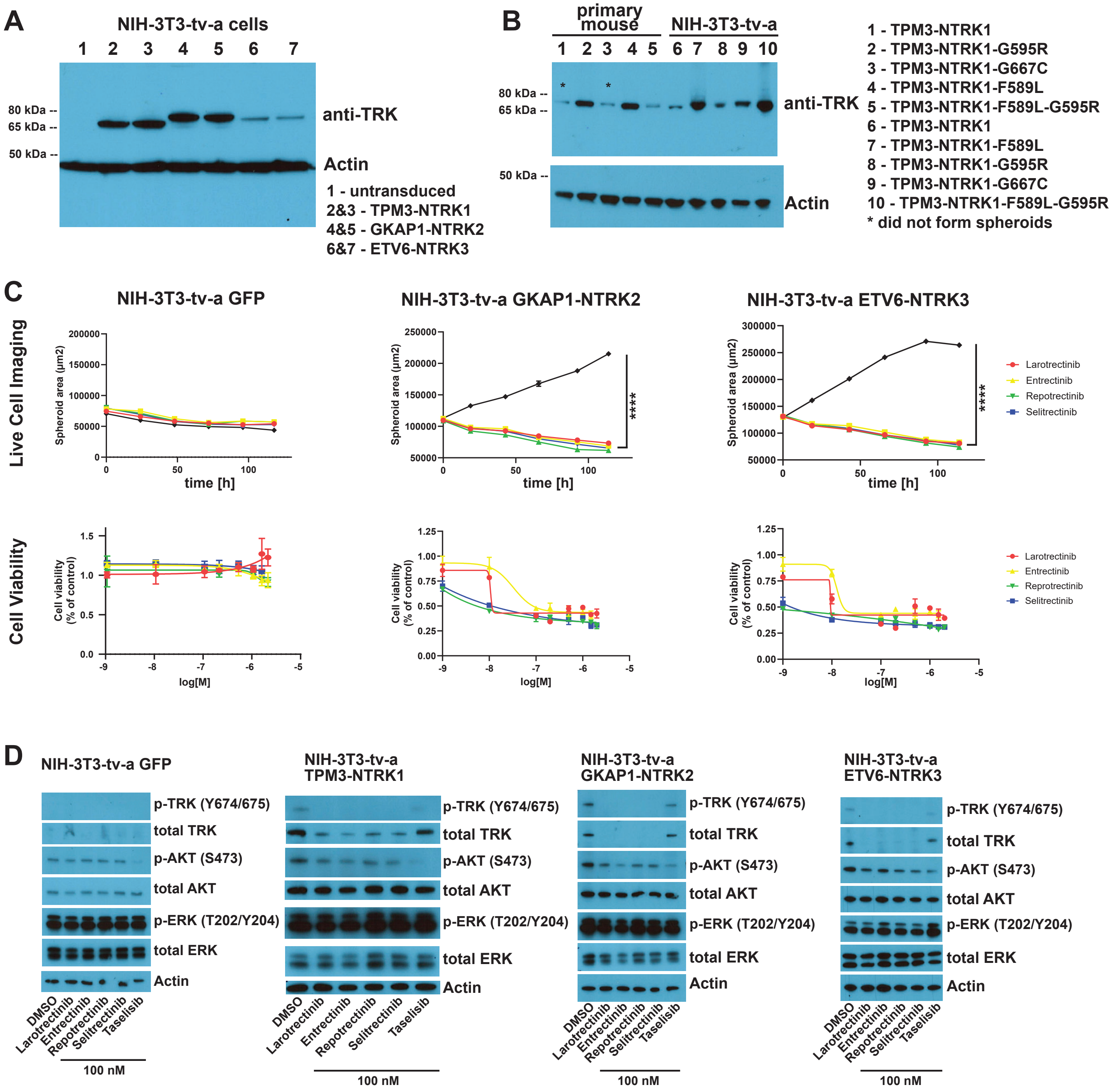

### Suppl. Figure S3E

**E**

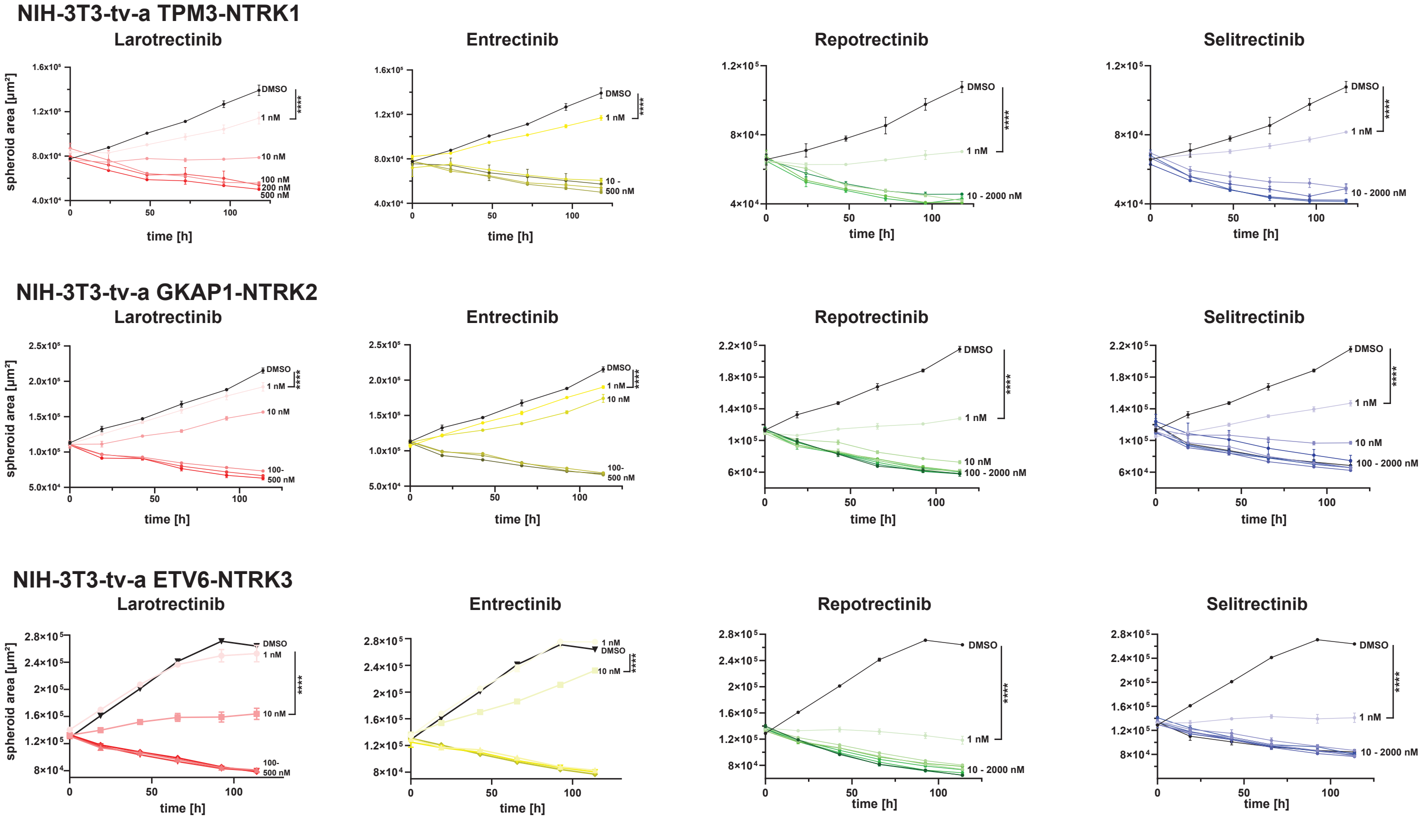

### Suppl. Figure S3F-I

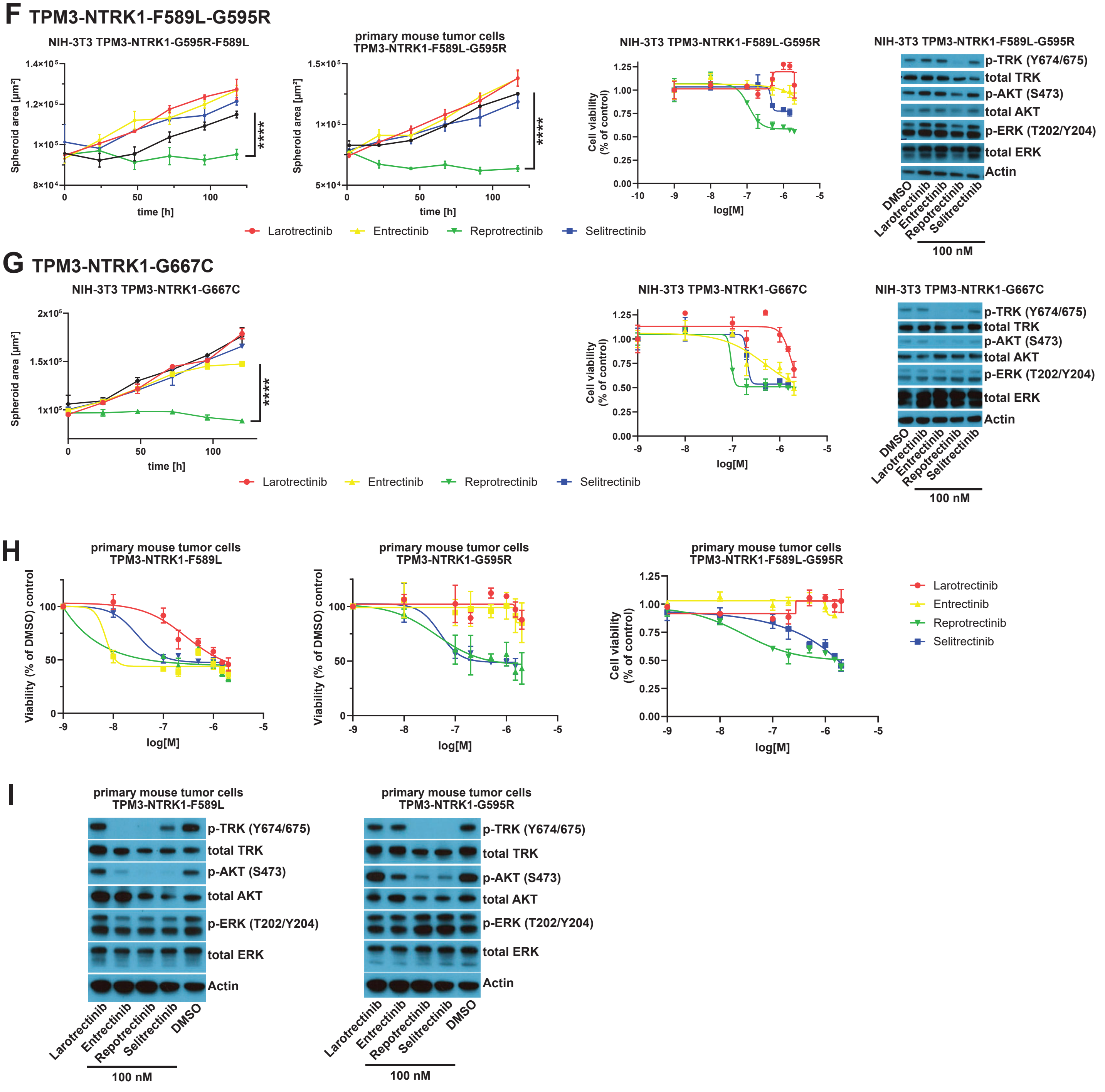

### Suppl. Figure S3J

## J

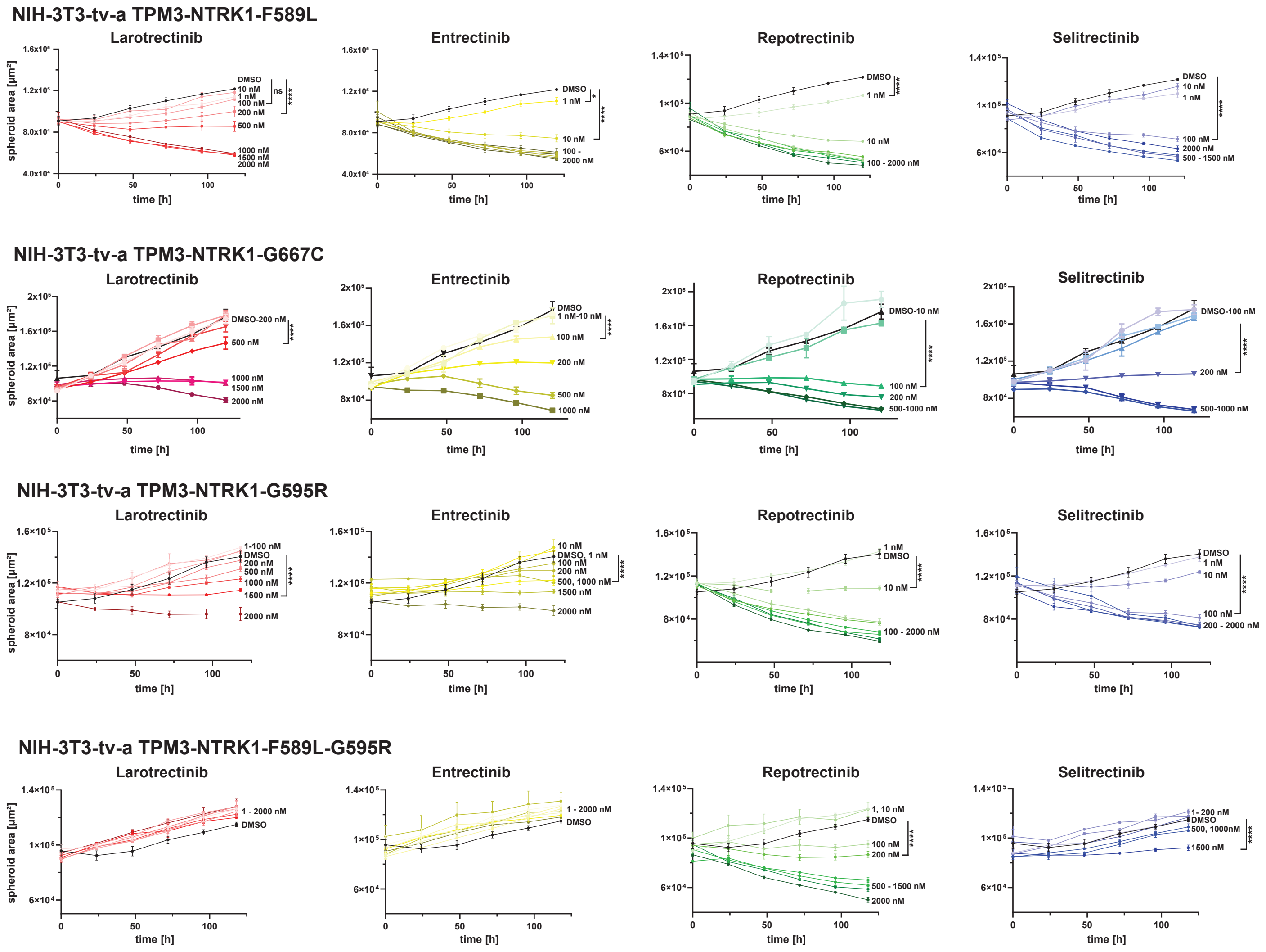

#### Suppl. Figure S4 E

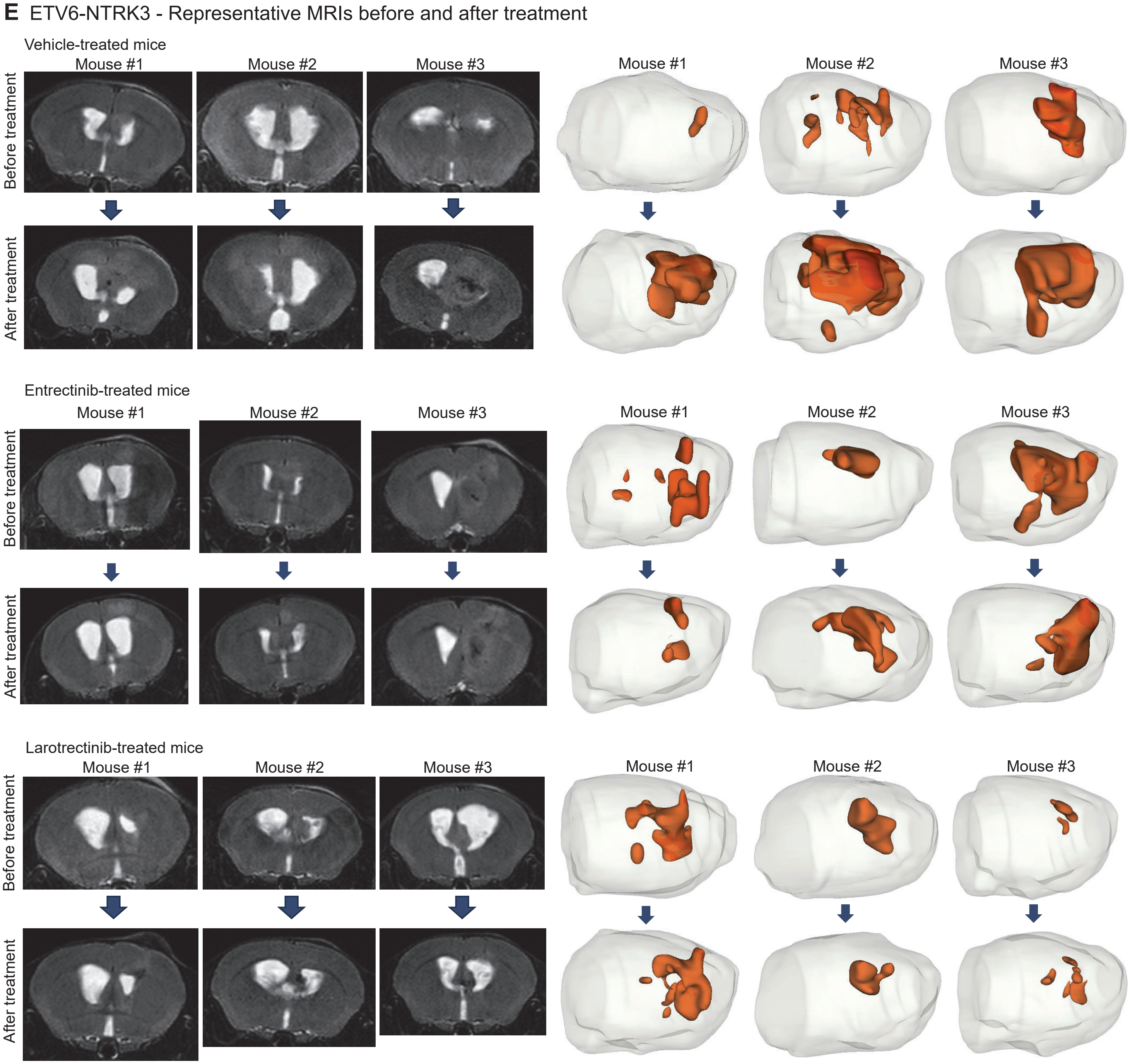

#### Suppl. Figure S4 A-C

**A** Recurrent tumors TPM3-NTRK1

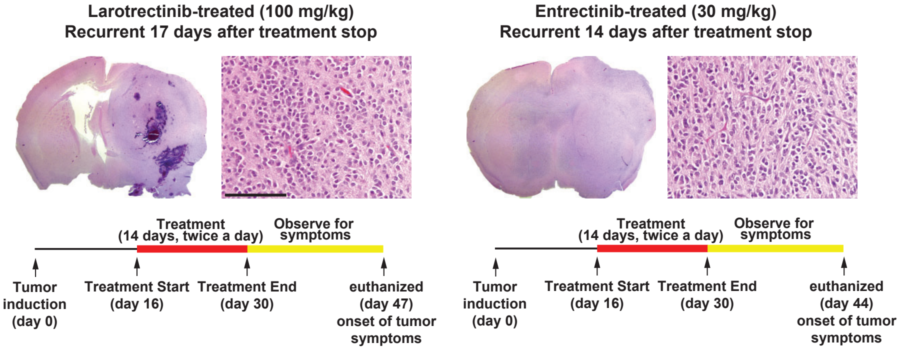


**B** Median survival of Vehicle- and TKI-treated NTRK fusion tumor-bearing mice (14 days of treatment)

| | Vehicle | Entrectinib | Larotrectinib |
|---|---|---|---|
| TPM3-NTRK1 | 38 | 44 | 46.5 |
| CHTOP-NTRK1 | 34 | 48 | 41 |
| GKAP1-NTRK2 | 36 | 40 | 45 |
| NACC2-NTRK2 | 42 | 50 | 56 |
| ETV6-NTRK3 | 41.5 | 53 | 57 |
| EML4-NTRK3 | 16 | 19 | 36 |

**C** TPM3-NTRK1 14 days of treatment

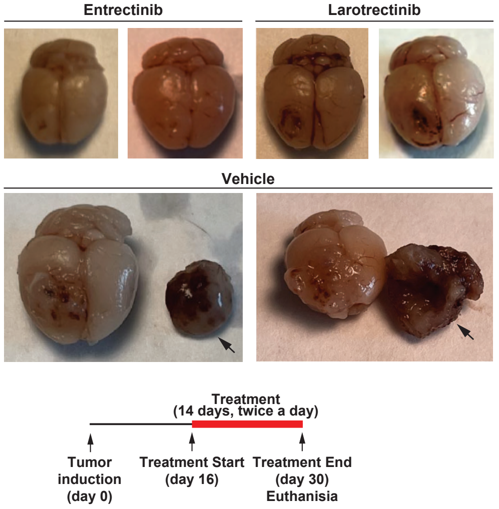

#### Suppl. Figure S4 D

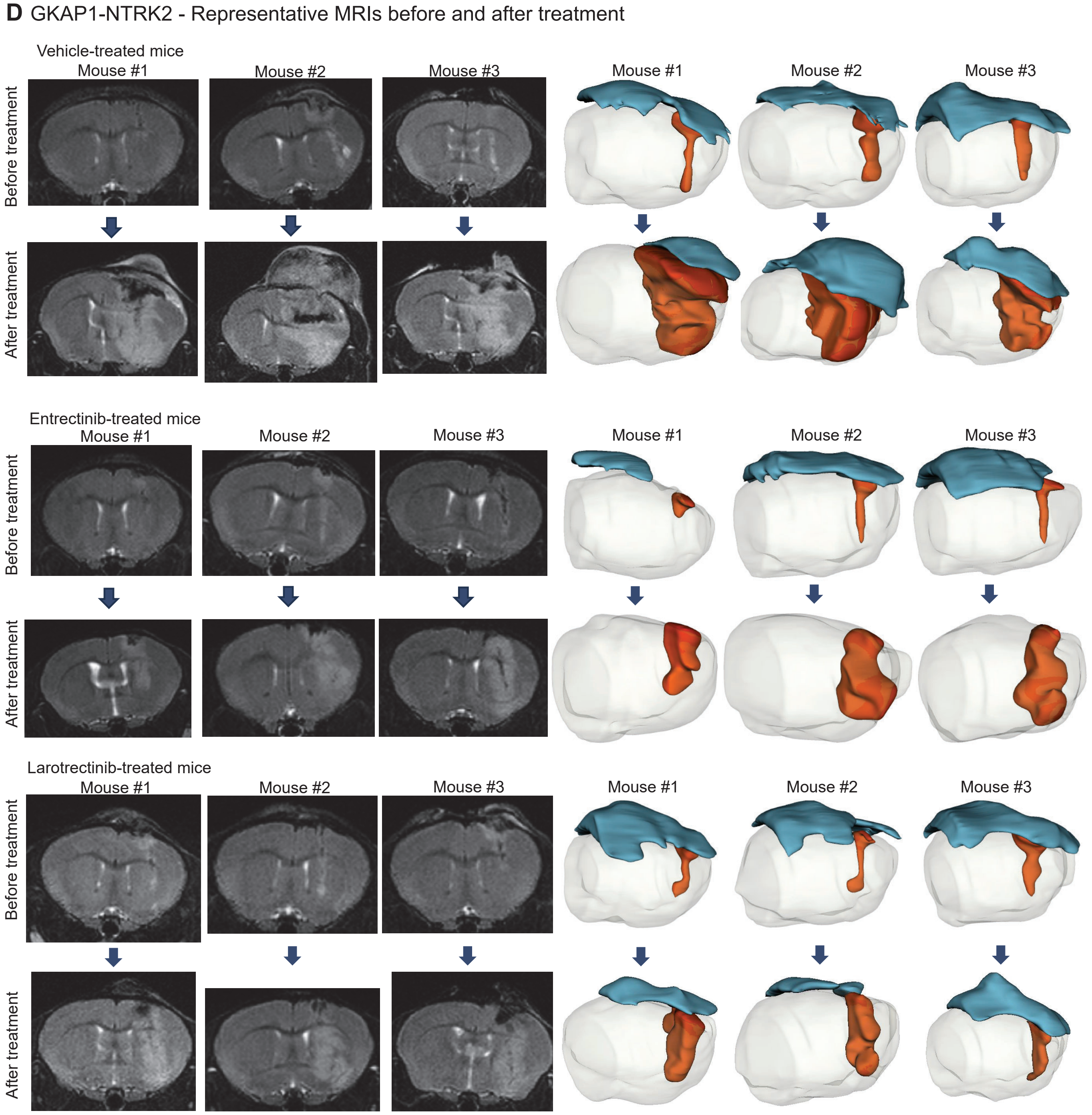

#### Suppl. Figure S4 F

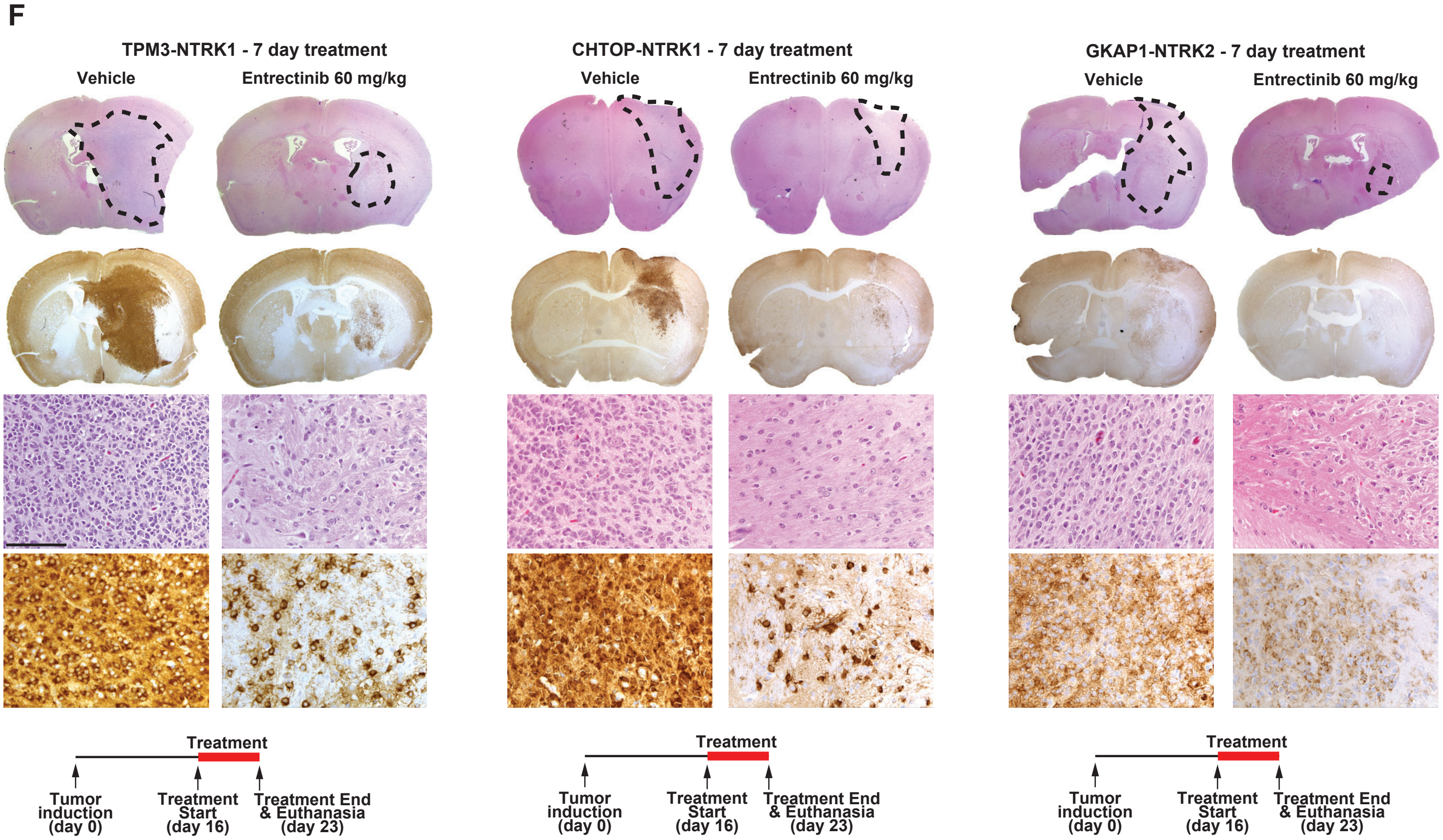

### Suppl. Figure S5A-B

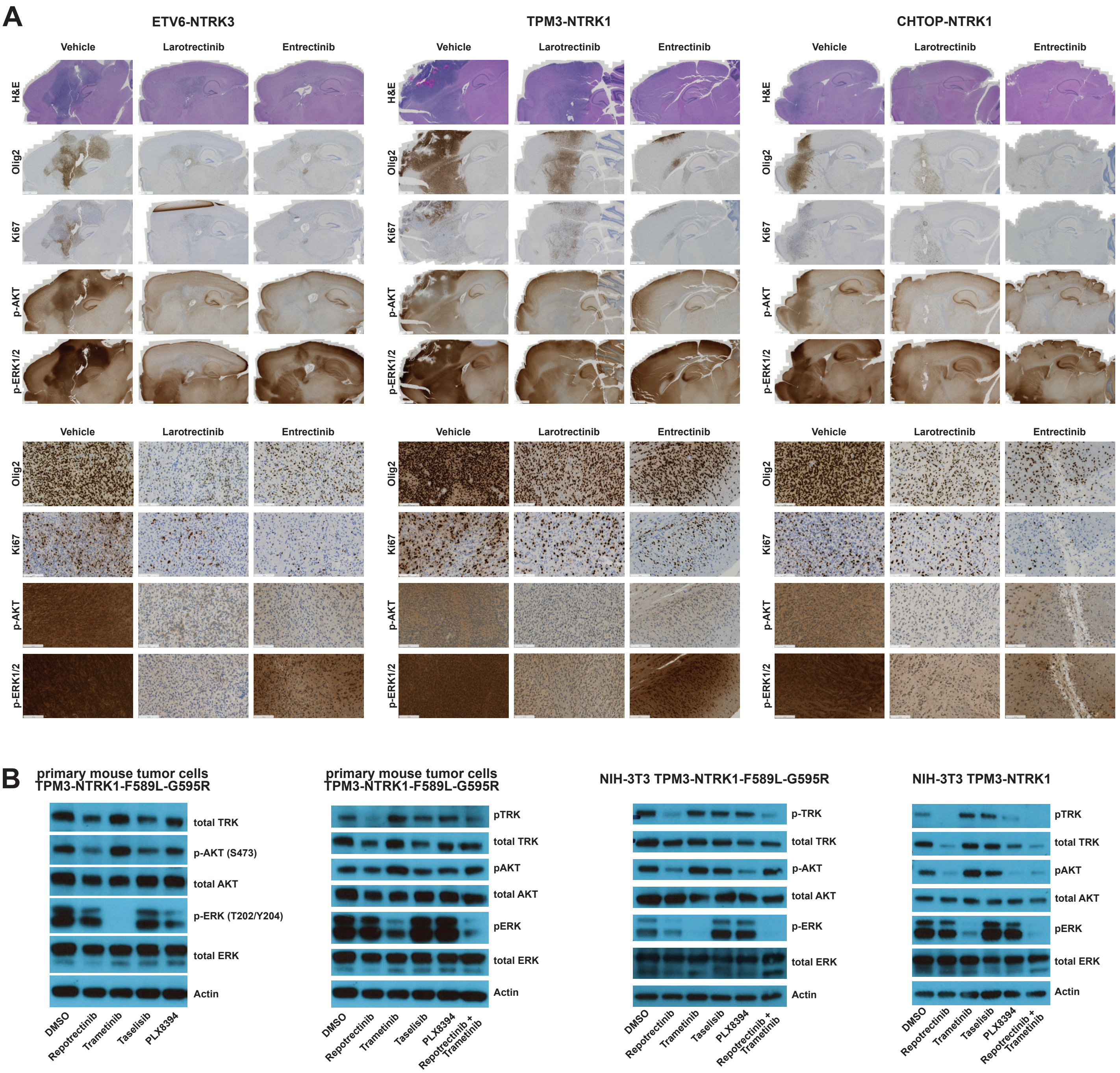

**Suppl. Figure legends:**

**Suppl. Figure S1: Expression of NTRK gene fusions in Nestin-positive cells induces the formation of experimental mouse gliomas.** A) Schematic representations of the three different NTRK receptors and their respective gene fusions, as well as fusion break points. C1/C1 = cysteine clusters, LRR = leucine-rich regions, Ig1-2 = immunoglobulin-like motifs, TM = transmembrane domain. B) Western Blot of DF1 cells transfected with RCAS plasmids expressing the different NTRK fusion genes. C) Overview table listing tumor penetrance and median survival of N/tv-a Cdkn2a wild type mice intracranially injected with RCAS-NTRK-fusion-expressing DF1 cells at different ages. D) Representative H&E images of tumors arising in N/tv-a Cdkn2a wild type mice intracranially injected with RCAS-NTRK-fusion-expressing DF1 cells at different ages. Black dotted lines indicate tumor margins. E) Representative images of H&E stainings of NTRK fusion tumors arising from the third ventricles of N/tv-a Cdkn2a wild type mice. F) Overview table listing tumor penetrance and median survival of N/tv-a Cdkn2a null mice intracranially injected with RCAS-NTRK-fusion-expressing DF1 cells at different ages. G) Kaplan-Meier curve showing symptom-free survival of N/tv-a Cdkn2a null mice intracranially injected as adults (at 5-7 weeks of age) with RCAS-NTRK fusion-expressing DF1 cells. H) Representative H&E images of tumors arising in N/tv-a Cdkn2a null mice intracranially injected with RCAS-NTRK-fusion-expressing DF1 cells at different ages. Black dotted lines indicate tumor margins. I) Kaplan-Meier curve showing symptom-free survival of N/tv-a Cdkn2a null mice intracranially injected to express either ETV6-NTRK3 (short variant) or ETV6-NTRK3-v2 (variant with longer TRK kinase domain). J) Representative images of IHC stainings for the presence of the glial markers Olig2 and GFAP in mouse gliomas derived by the expression of TPM3-NTRK1 and CHTOP-NTRK1 in N/tv-a Cdkn2a null mice. K) Overview table listing tumor penetrance and median survival of N/tv-a Cdkn2a wild type mice intracranially injected with RCAS-NTRK-fusion and RCAS-shPten-expressing DF1 cells. L) Representative images of H&E and IHC stainings (TRK kinase domain, Olig2, GFAP) of the extraaxial, the intracranial, and the extra-cranial tumor components of a glioma induced by the intracranial expression of RCAS-TPM3-NTRK1 in adult N/tv-a Cdkn2a null mice. M) Representative MRI images (T2w, T1w pre-contrast, T1w post-contrast) of a glioma induced by the intracranial expression of TPM3-NTRK1 in adult N/tv-a Cdkn2a null mice. N-O) Overview table listing tumor penetrance and median survival (N) and representative H&E images (O) of N/tv-a Cdkn2a null mice i.p. injected with RCAS-TPM3-NTRK1-expressing DF1 cells. P) Overview table listing tumor penetrance and median survival of N/tv-a Cdkn2a wild type and Cdkn2a null mice intracranially injected with DF1 cells expressing the different point mutant variants of RCAS-TPM3-NTRK1. Q-R) Kaplan Meier curves showing symptom-free survival of N/tv-a Cdkn2a wild type (Q) and Cdkn2a null (R) mice intracranially injected with DF1 cells expressing the different point mutant variants of RCAS-TPM3-NTRK1. S-T) Representative H&E images of N/tv-a Cdkn2a null (S) and Cdkn2a wild type (T) mice intracranially injected with DF1 cells expressing the different point mutant variants of RCAS-TPM3-NTRK1. Analysis was done using Log-rank (Mantel-Cox) test (I,Q,R). Scale bars indicate 100 µm.

**Suppl. Figure S2: NTRK fusion-driven experimental mouse gliomas show activation of the PI3K-AKT-S6 and RAF-MEK-ERK pathways.** Representative IHC stainings (TRK (TRK kinase domain), Olig2, GFAP, Ki-67, phospho-ERK1/2 (Thr202/Thr185), phospho-AKT (Ser473), and phospho-S6 (Ser 240/244)) of mouse gliomas induced by the intracranial injection of RCAS-NTRK fusion-expressing DF1 cells into N/tv-a Cdkn2a null mice. A) TPM3-NTRK1 B) CHTOP-NTRK1 C) GKAP1-NTRK2 D) QKI-NTRK2 E) NACC2-NTRK2 F) ETV6-NTRK3 G) BTBD1-NTRK3 H) EML4-NTRK3. Scale bar indicates 100 µm.

**Suppl. Figure S3: NTRK fusion-driven cells respond to TKI treatment in vitro.** A) Western Blot stained for TRK (Kinase domain) and Actin, showing the expression of TPM3-NTRK1, GKAP1-NTRK2, and ETV6-NTRK3 in NIH-3T3-tv-a cells transduced with RCAS viruses. B) Western Blot stained for TRK (Kinase domain) and Actin showing the expression of different TPM3-NTRK1 point mutant versions in primary mouse tumor cells and in vitro transduced NIH-3T3-tv-a cells. C) Growth curves (upper panels) and dose response curves (lower panels) of NIH-3T3-tv-a cell spheroids expressing either GFP (left panels), GKAP1-NTRK2 (middle panels), or ETV6-NTRK3 (right panels). Growth curves show treatment of spheroids with either DMSO (0.1%) or TKIs (Larotrectinib, Entrectinib, Selitrectinib, Repotrectinib; 100 nM each). For dose response curves cell spheroids were treated with different TKI concentrations (1, 10, 100, 200, 500 nM, 1 µM, 1.5 µM, or 2 µM) and viability is shown relative to DMSO-treated cells. D) Representative Western Blots of NIH-3T3-tva cells expressing either GFP (left panel), TPM3-NTRK1 (left middle panel), GKAP1-NTRK2 (right middle panel), or ETV6-NTRK3 (right panel). Cells were treated with either DMSO (0.1%), TKIs (Larotrectinib, Entrectinib, Repotrectinib, or Selitrectinib), or the PI3K inhibitor Taselisib (100 nM for all). E) Growth curves of NIH-3T3-tv-a cell spheroids expressing either TPM3-NTRK1 (upper panels), GKAP1-NTRK2 (middle panels), or ETV6-NTRK3 (lower panels). Growth curves show growth of cell spheroids upon treatment with different concentrations of TKIs (Larotrectinib, Entrectinib, Repotrectinib, Selitrectinib) at indicated concentrations. F) Effect of the TPM3-NTRK1-F589L-G595R kinase domain mutation on efficacy of TKIs. Left and middle left panels: Growth of TPM3-NTRK1-F589L-G595R-expressing NIH-3T3-tv-a (left panel) and primary mouse tumor cell (middle left panel) spheroids upon treatment with TKIs (Larotrectinib, Entrectinib, Repotrectinib, Selitrectinib, 100 nM each). Middle right panel: Dose response curve of TPM3-NTRK1-F589L-G595R-expressing NIH-3T3-tv-a cell spheroids treated with different TKI concentrations (1, 10, 100, 200, 500 nM, 1 µM, 1.5 µM, or 2 µM). Right panel: Representative Western Blots of TPM3-NTRK1-F589L-G595R-expressing NIH-3T3-tva cells treated with TKIs (Larotrectinib, Entrectinib, Repotrectinib, Selitrectinib, 100 nM each). G) Effect of the TPM3-NTRK1-G667C kinase domain mutation on efficacy of TKIs. Left panel: Growth of TPM3-NTRK1-G667C-expressing NIH-3T3-tv-a cell spheroids upon treatment with TKIs (Larotrectinib, Entrectinib, Repotrectinib, Selitrectinib, 100 nM each). Middle right panel: Dose response curve of TPM3-NTRK1-G667C-expressing NIH-3T3-tv-a cell spheroids treated with different TKI concentrations (1, 10, 100, 200, 500 nM, 1 µM, 1.5 µM, or 2 µM). Right panel: Representative Western Blots of TPM3-NTRK1-G667C-expressing NIH-3T3-tva cells treated with TKIs (Larotrectinib, Entrectinib, Repotrectinib, Selitrectinib, 100 nM each). H) Dose response curves of primary mouse tumor cell spheroids expressing either TPM3-NTRK1-F589L (left panel), TPM3-NTRK1-G595R (middle panel), or TPM3-NTRK1-F589L-G595R (right panel). Spheroids were treated with different TKI concentrations (1, 10, 100, 200, 500 nM, 1 µM, 1.5 µM, or 2 µM). I) Representative Western Blots of primary mouse tumor cells expressing either TPM3-NTRK1-F589L (left panel) or TPM3-NTRK1-G595R (right panel). Cells were treated with TKIs (Larotrectinib, Entrectinib, Repotrectinib, Selitrectinib, 100 nM each). J) Growth curves of NIH-

3T3-tv-a cell spheroids expressing either TPM3-NTRK1-F589L (upper panels), TPM3-NTRK1-G667C (upper middle panels), TPM3-NTRK1-G595R (lower middle panels), or TPM3-NTRK1-F589L-G595R (lower panels). Growth curves show growth of cell spheroids upon treatment with different concentrations of TKIs (Larotrectinib, Entrectinib, Repotrectinib, Selitrectinib) at indicated concentrations. Analysis was done using ordinary two-way ANOVA test (C,E,F,G,J).

**Suppl. Figure S4: TKI treatment significantly prolongs the survival of NTRK gene fusion-driven mouse gliomas.** A) Representative images of recurrent tumors. B) Overview table showing median survival of mice harboring gliomas induced by the RCAS-mediated intracranial expression of the different NTRK fusions. Mice were treated with either DMSO, Entrectinib (30 mg/kg), or Larotrectinib (100 mg/kg) via i.p. injection b.i.d. for 14 days. C) Representative images of whole brains (and extracranial tumors) of N/tv-a Cdkn2a null mice harboring gliomas induced by the intracranial expression of RCAS-TPM3-NTRK1 after 14 days of treatment with either DMSO, Entrectinib (30 mg/kg), or Larotrectinib (100 mg/kg) via i.p. injection b.i.d. Mice were euthanized 1 hour after the last treatment. D-E) Representative cranial MRIs and 3D tumor brain reconstructions generated from mice harboring either (D) RCAS-GKAP1-NTRK2 or (E) RCAS-ETV6-NTRK3-driven gliomas before and after treatment (14 days, i.p., b.i.d.) with either Vehicle (DMSO), Entrectinib (30 mg/kg), or Larotrectinib (100 mg/kg). F) Representative H&E and IHC (for TRK kinase domain) stainings of intracranial RCAS-TPM3-NTRK1, RCAS-CHTOP-NTRK1, or RCAS-GKAP1-NTRK2-driven mouse gliomas after 7 days of treatment (i.p., b.i.d.) with either Vehicle (DMSO) or Entrectinib (60 mg/kg).

**Suppl. Figure S5: Combination therapy of TRK and MEK inhibition is superior to TRK inhibition alone against NTRK fusion-driven cells.** A) Representative H&E and IHC stainings (Olig2, Ki-67, phospho-AKT (Ser473); phospho-ERK1/2 (Thr202/Thr185)) of *ETV6-NTRK3, TPM3-NTRK1, and CHTOP-NTRK1*-mouse gliomas (Vehicle or 14 days of Larotrectinib (100 mg/kg) or Entrectinib (30 mg/kg) treatment). Scale bars indicate 1 mm (whole brain slides) or 100 µm (high magnification panels). B) Representative Western Blot of primary TPM3-NTRK1-F589L-G595R mouse tumor cells or NIH-3T3 cells expressing either TPM3-NTRK1 or TPM3-NTRK1-F589L-G595R treated with either DMSO (0.1%; Vehicle), Repotrectinib (TKI), Trametinib (MEKi), Taselisib (PI3Ki), PLX8394 (BRAFi), or a combination of Repotrectinib plus Trametinib (TKI + MEKi). All inhibitors were used at 100 nM except Trametinib at 10 nM.

#### Supplemental experimental procedures

##### Plasmid generation
Primers used for plasmid generation are listed in Supplemental Table S1A. Additional plasmids used in this study are listed in Supplemental Table S1C.

##### Transfection of RCAS viruses
Chicken fibroblast (DF1) cells were maintained with 10% fetal bovine serum (FBS) in Dulbecco's modified Eagle medium (DMEM) (including 1% Penicillin/Streptomycin) at 39 degrees Celsius. DF-1 cells were transfected with the indicated RCAS plasmid using X-tremeGENE 9 DNA transfection reagent (Roche) according to the manufacturer's protocol. RCAS transgene expression was confirmed via western blot analysis.

##### Generation of NTRK fusion-expressing NIH-3T3 cells
Viral supernatant from DF1 cultures was sterile-filtered and added to Tv-a-expressing NIH-3T3 cells [4] every eight hours. NIH-3T3-tv-a cells were maintained with 10% Newborn CALF serum (Thermo Fisher 26010074) in DMEM (including 1% Penicillin/Streptomycin) at 37 degrees Celsius. RCAS transgene expression was confirmed via western blot analysis.

##### Culturing of primary mouse tumor cells
Primary mouse tumors generated by flank injection of RCAS-producing DF1 cells into Nestin/tv-a Cdkn2a null mice were manually dissected using scalpels and maintained with 10% FBS in DMEM (including 1% Penicillin/Streptomycin) at 37 degrees Celsius. RCAS transgene expression was confirmed via western blot analysis.

##### Spheroid assay
NIH-3T3-tv-a cells (expressing either GFP, TPM3-NTRK1 (or mutants thereof), GKAP1-NTRK2, or ETV6-NTRK3) or primary mouse tumor cells were seeded at $4x10^3$ cells per well in a 96-well ultra-low attachment plates (Corning 7007) in DMEM with 10% FBS and spun down at 1250 rpm for 10 minutes. After 2 days, cells were treated with DMSO or TKIs at indicated concentrations. Growth of spheroids was monitored using live cell imaging every 24 hours for 5 days in the Incucyte ZOOM system (Essen). Average phase object area ($mm^2$) was used for analysis. Viability was measured using the CellTiter-Glo® 2.0 Cell Viability Assay (Promega G9241) according to the manufacturer's instructions. Luminescence was detected using a Veritas Microplate Luminometer.

##### Magnetic resonance imaging (MRI)
Tumor size and progression were evaluated using the preclinical 7 Tesla MR scanner (MR Solutions, DRYMAG7.0T) located in the FHCC imaging suite. Following anesthetic induction with 2% isoflurane, each mouse was placed on a bed and respiration was monitored via pneumatic pillow (SA Instruments, ERT gating module). Mouse body temperature was maintained by warm air circulation within the arm. Each procedure lasted approximately 10-30 minutes. Images were acquired using a mouse brain RF coil. Techniques used are Fast Spin Echo (FSE) T2 weighted scans (FSE T2w (axial) TE=75, 15 slices at 0.5mm thickness, FOV 25, 1 average) and for contrast enhanced imaging FSE T1 weighted scan pre- and post-contrast injection (FSE T1w (axial) TE=11, 15 slices at 0.5mm thickness, FOV 25, 2 averages). For contrast enhanced imaging 1 mL Gadobenate (MultiHance 529mg/ml, 0.5M) was diluted in 9 mL saline and mice were dosed with

4 ul/g of solution. Following imaging, each mouse was allowed to recover from anesthesia in a warm recovery cage. Tumor volumes based on MRI images were calculated using the software 3D slicer and graphed with GraphPad Prism.

**H&E Staining, Immunohistochemistry stainings of FFPE mouse tissues**

For routine tumor histomorphology, mouse brains and peripheral tumors were formalin-fixed and paraffin-embedded, sectioned, and stained with H&E as described previously [1, 3]. For stainings in Figure 2 and Suppl. Figure 2, mouse brains were cut along the sagittal plane into 4 pieces, flash frozen in liquid nitrogen, and subsequently thawed in formalin and subsequently processed as described above. Immunohistochemical (IHC) staining of mouse brains was performed on the DISCOVERY XT platform (Ventana Medical Systems, Inc., Tucson, U.S.A) using the Discovery DAB Map Detection Kit according to standard protocols as described previously [2]. For a list of antibodies used see Suppl. Table S1B. IHC images were quantified using HistoQuant software.

**Western Blot Analysis**

Cells were cultured, lysed, and processed for western blotting by standard methods. Proteins were resolved by SDS/PAGE (NuPAGE 10% Bis/Tris; LifeTech) according to XCell Sure Lock Mini-Cell guidelines, blocked with 5% milk/TBST and probed with specified antibodies overnight at 4°C in 5% BSA/TBST. After three TBST rinses, species-specific secondary antibodies were added in 5% milk/TBST. Blots were rinsed three times with TBST before being developed with Amersham ECL Western Blotting Detection Reagents (GE Healthcare). For a list of antibodies used see Suppl. Table S1B.

**Chemicals**

Larotrectinib (LOXO-101, NSC 785570), Entrectinib (NMS-E628, NSC 774769), Repotrectinib (TPX-0005, NSC 800522), and Selitrectinib (LOXO-195, NSC 809970) were provided through the National Institutes of Health (NIH) Developmental Therapeutics Program. Trametinib (GSK1120212, Cat. S2673), Taselisib (GDC 0032, Cat. S7103), and PLX8394 (Cat. S7965) were purchased from Selleckchem.

**Determining TKI doses for in vivo treatment**

Mice were treated by i.p. injection of drug twice-a-day (b.i.d.). To assess the optimal in vivo inhibitor concentrations for treatment, we first treated non-tumor mice with different doses of either Entrectinib (20, 30, 40, 60 mg/kg, two mice per group) or Larotrectinib (46, 92, 138 mg/kg, three mice per group) for several days to assess general drug toxicity. One mouse treated with 20 mg/kg Entrectinib died during treatment, while one mouse treated with 92 mg/kg Larotrectinib died two days after the last treatment. In general, we observed that mice treated with either 92 or 138 mg/kg Larotrectinib developed distended abdomens due to fluid retention that receded several days after treatment discontinuation. We first treated several mice with intracranial gliomas with Larotrectinib (60 or 100 mg/kg), Entrectinib (30 or 60 mg/kg), or Vehicle for 7 days and subsequently analyzed the treated tumors by histology. We observed a greater tumor response at 100 mg/kg Larotrectinib compared to 60 mg/kg and no toxicity-related deaths in either cohort. Similarly, we observed a greater response at 60 mg/kg Entrectinib compared to 30 mg/kg, however, observed several toxicity-related deaths at 60 mg/kg. Based on these results, we proceeded with a dosing strategy of 100 mg/kg Larotrectinib and 30 mg/kg Entrectinib.

**Treatment scheduling for different fusion tumors**

The first MRI was performed 15-25 days post-injection (for exact times see below). Treatment started one day after the MRI.
TPM3-NTRK1 (15 days post-injection), CHTOP-NTRK1 (15 days post-injection); GKAP1-NTRK2 (15 days post-injection); NACC2-NTRK2 (18 days post-injection); ETV6-NTRK3 (25 days post-injection)
